# Mitochondrial control of amino acid catabolism by a fasting-inducible mitochondrial carrier

**DOI:** 10.64898/2026.05.26.727879

**Authors:** Satoshi Oikawa, Tadashi Yamamuro, Hiroshi Nishida, Daisuke Katoh, Dandan Wang, Masanori Fujimoto, Shingo Kajimura

## Abstract

Metabolic adaptation to nutrient deprivation requires coordinated control of mitochondrial anaplerosis and cataplerosis; however, how metabolite flux across the mitochondrial membrane is regulated during fasting remains less defined. Here, we report SLC25A34 as a fasting-inducible mitochondrial carrier that is highly expressed in oxidative skeletal muscle. Using bacterial reconstitution, proteo-liposomes, and tracer studies, we showed that SLC25A34 mediates the import of phosphoenolpyruvate (PEP) into the mitochondrial matrix. Loss of SLC25A34 impaired glutamine-supported anaplerosis under nutrient-deprived conditions, while glucose and pyruvate utilization remained largely intact. Muscle-specific deletion of *Slc25a34* resulted in reduced fasting-induced amino acid catabolism and the accumulation of amino acids, leading to activation of mTORC1 signaling even under fasted conditions. Consequently, SLC25A34-deficient soleus muscle exhibited hypertrophy and myopathic features, accompanied by mTORC1-dependent increase in protein synthesis. Together, these results highlight a unique biological role for the inducible mitochondrial carrier SLC25A34, which couples PEP import to amino acid catabolism and proteostasis to preserve skeletal muscle integrity in response to metabolic stress.

**Teaser:** Mitochondrial metabolite transport is tailored to meet muscle-specific needs for utilizing amino acids during fasting.

## INTRODUCTION

Mitochondria serve far broader roles than ATP production through integrating nutrient availability with fuel selection to support adaptation to physiological and pathological stress (*1–3*). Under conditions of limited glucose availability, such as prolonged fasting, peripheral tissues, including skeletal muscle and brown adipose tissue (BAT), shift away from carbohydrate use and increase the oxidation of fatty acids released by adipose tissue lipolysis (*4–6*). Skeletal muscle also undergoes proteolysis, supplying amino acids for hepatic gluconeogenesis, while sustaining its own bioenergetic needs to preserve glucose for the brain, an obligate glucose-dependent organ (*7, 8*). Similarly, endurance exercise requires flexible substrate utilization in muscle to balance carbohydrate sparing with sustained energy production (*4, 9, 10*). In turn, under pathological conditions, such as heart failure, the failing heart shifts away from fatty acid oxidation and relies on glucose metabolism (*11*).

In this context, a key regulator is the balance between anaplerosis and cataplerosis of the tricarboxylic acid (TCA) cycle. Anaplerosis replenishes TCA cycle intermediates to maintain oxidative capacity, whereas cataplerosis enables the exit of carbon from the cycle to support biosynthesis, signaling, and redox homeostasis, as well as to prevent excessive accumulation of intermediates in the matrix (*12–15*). For example, acetyl-CoA derived from pyruvate or fatty acid oxidation condenses with oxaloacetate (OAA) to form citrate; glutamine contributes anaplerotic carbon via conversion to α-ketoglutarate (αKG); and branched-chain amino acids feed the cycle through acetyl-CoA (Leu, Ile) or succinyl-CoA (Val). On the other hand, cataplerotic flux, such as the export of citrate, supports lipid synthesis and acetylation pathways (*16, 17*). Additionally, the export of malate and OAA maintains redox balance and serves as substrate shuttles (*13–15*). Despite the importance of this balance, the regulatory mechanisms that coordinate anaplerosis and cataplerosis during metabolic adaptation remain less explored.

A knowledge barrier has been the limited understanding of the molecular determinants controlling metabolite flux across the inner mitochondrial membrane (IMM). Since the IMM is impermeable to most metabolites, anaplerosis and cataplerosis depend on dedicated metabolite carriers localized to the IMM (*18*). For instance, the mitochondrial pyruvate carrier (MPC) is essential for importing pyruvate into the mitochondrial matrix (*19, 20*), while SLC25A1 (CIC) and SLC25A10 (DIC) facilitate the transport of citrate and dicarboxylic acids (e.g., succinate), respectively (*21–24*). SLC25A11 (OGC) and SLC25A12/13 (AGC1/2) mediate the malate-aspartate shuttle that supports NADH shuttling across the IMM (*25–28*). Of note, many of these IMM carrier proteins are ubiquitously and constitutively expressed; however, some carriers are tissue-selective and uniquely regulated by external cues. An example is uncoupling protein 1 (UCP1, also known as SLC25A7 for H^+^ import), which is highly induced by cold exposure and the beta-adrenergic signaling pathway in brown and beige adipocytes (*29, 30*). Thus, it is likely that stress-inducible IMM carriers expressed in a tissue-selective manner serve as an additional regulatory layer of metabolic adaptation by modulating anaplerotic and cataplerotic fluxes depending on the cellular context.

The present study reports SLC25A34 as an example of such: SLC25A34 shares the highest sequence homology with SLC25A35 among the SLC25A family, which we recently identified as a mitochondrial carrier for phosphoenolpyruvate (PEP) (*31*). PEP is a unique metabolite from a view of bioenergetics, as PEP contains one of the highest energy phosphate bonds (-61.9 kJ mol ^-1^), higher than ATP (-30.5 kJ mol ^-1^), and has diverse biological roles, including glycolysis, gluconeogenesis, glyceroneogenesis, and insulin secretion (*32–35*). We found that SLC25A34 is highly expressed in skeletal muscle, particularly in the mitochondria-rich soleus muscle at a higher level than SLC25A35 (**Supplementary Figure 1A, B**). Notably, unlike SLC25A35, SLC25A34 expression is induced by metabolic stressors, including fasting and exercise. Hence, the present work investigated the biological role of SLC25A34 in regulating the balance of mitochondrial anaplerosis and cataplerosis in muscle in response to metabolic stress.

## RESULTS

### SLC25A34 transports phosphoenolpyruvate (PEP)

SLC25A34 belongs to the SLC25A protein family, many of which are expressed in the mitochondrial membrane (*24, 36*), and shares a predicted structural similarity with SLC25A35, exhibiting 63.7% amino acid sequence similarity. We first validated the cellular localization of the SLC25A34 protein by immunostaining, confirming its presence in the IMM (**Supplementary Figure 1C**). The predicted structure of SLC25A34, as generated by AlphaFold (*37*), suggests a highly similar overall architecture to SLC25A35, which consists of six transmembrane α-helices, a characteristic of SLC25 carriers (**Figure 1A**). Our recent study in SLC25A35 identified critical amino acids responsible for substrate recognition, including Y72, Q73, M76, R80, Y124, K127, R175, R276, and H280 (*31*). Notably, these residues are well-conserved between SLC25A34 and SLC25A35 (**Figures 1B, C**), suggesting that they share common substrates.

**Figure 1.**
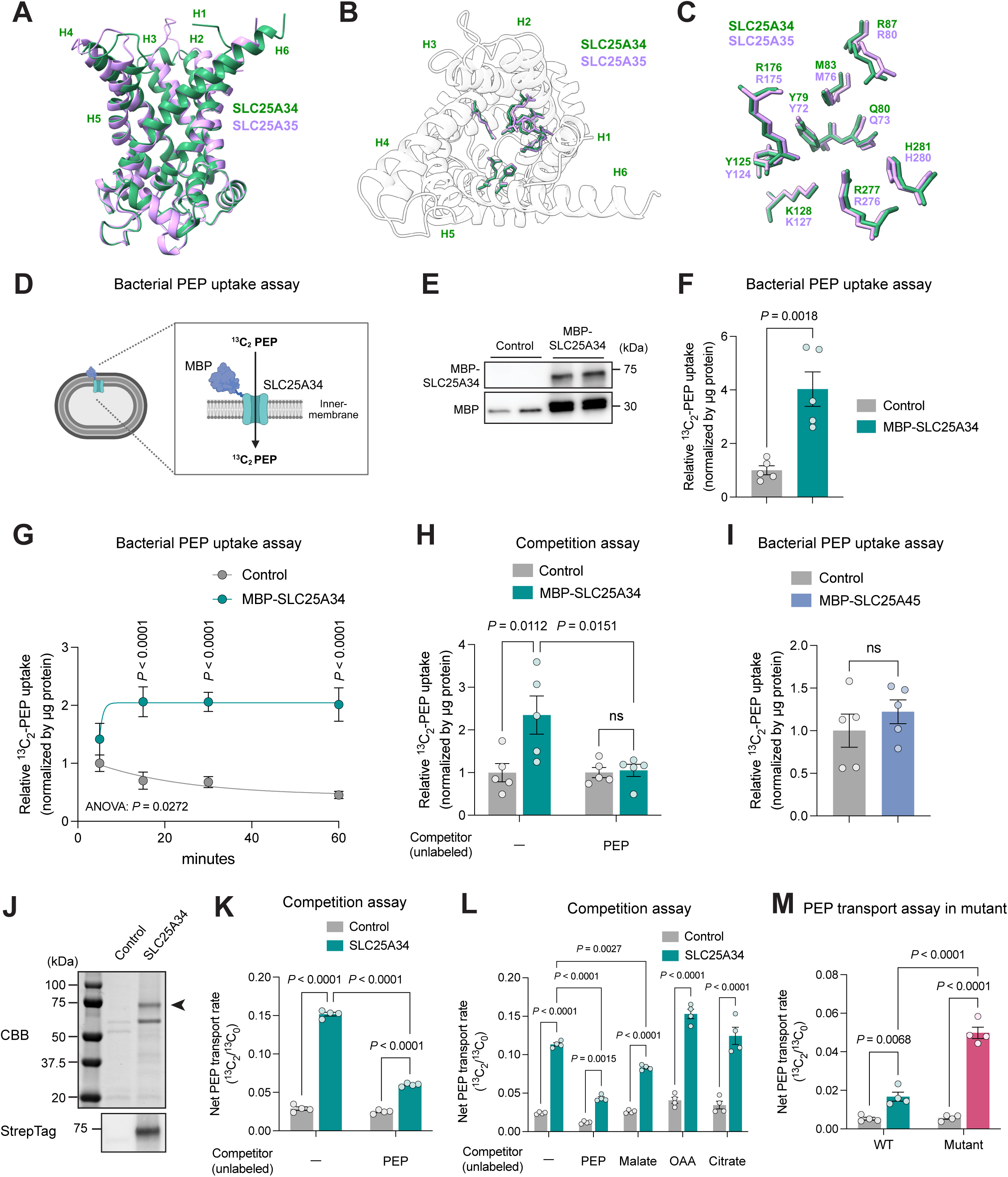
SLC25A34 transports phosphoenolpyruvate (PEP). **A.** Side view of the predicted SLC25A34 structure (green) aligned with the predicted SLC25A35 structure (purple), created using UCSF ChimeraX. **B.** Top view of the aligned SLC25A34 (green) and SLC25A35 (purple) structures. Key amino acid residues involved in phosphoenolpyruvate (PEP) binding within the central channel-like cavity were highlighted. **C.** Close-up of the key amino acid residues of SLC25A34 (green) and SLC25A35 (purple) in the central channel-like cavity. **D.** Schematic of PEP uptake assay with bacteria. **E.** Immunoblot of MBP-SLC25A34 protein in *E. coli*. Note that *E. coli* expresses the endogenous MBP. **F.** PEP uptake in control and MBP-SLC25A34-expressing *E. coli*. N = 5 per group. **G.** Time course of PEP uptake in control and MBP-SLC25A34-expressing *E. coli*. N = 5 per group. **H.** Competition assay. An excess amount of unlabeled PEP (50 mM) was added to the bacterial uptake assay. N = 5. **I.** PEP uptake in control or MBP-SLC25A45-expressing *E. coli*. N = 5 per group. **J.** CBB staining and immunoblot of affinity-purified SLC25A34 protein. Arrowhead indicates the MBP-SLC25A34-TwinStrep protein. **K.** PEP transport assay in proteo-liposomes reconstituted with purified SLC25A34 or control-eluates, with or without excess unlabeled PEP (50 mM). n = 4 per group. **L.** Competition assay. An excess amount of indicated metabolites (50 mM) was added to the assay. N = 4. **M.** PEP transport assay in proteo-liposomes reconstituted with wild-type or mutant (Y125A/R176A/R277A) SLC25A34, or control-eluates. N = 4 per group. Bars represent mean ± s.e.m. *P* values were calculated by unpaired *t*-test (F and I) and two-way ANOVA with Tukey’s multiple comparisons test (G, H, K, L and M).

To test the hypothesis that SLC25A34 also transports PEP, we employed a bacterial reconstitution system in which human SLC25A34 protein was fused to maltose-binding protein (MBP) at the N-terminus in the *E. coli* strain C43 (DE3). This system has reduced Lon and OmpT protease activity, making it suitable for overexpressing membrane proteins and ensuring their correct orientation in the bacterial inner membrane (*38*) (**Figure 1D**). Although a previous study reported that the human SLC25A34 protein was not expressed in *E. coli* (*39*), the recombinant MBP-tagged SLC25A34 protein was successfully expressed in this system (**Figure 1E**). Note that *E. coli* expresses the endogenous MBP. Subsequently, the control *E. coli* expressing the endogenous MBP alone, as well as *E. coli* expressing MBP-SLC25A34, were incubated with ^13^C-labeled PEP, washed, and subjected to liquid chromatography-mass spectrometry (LC-MS) to evaluate their PEP import activity. We found that ^13^C-PEP uptake was significantly higher in *E. coli* expressing MBP-SLC25A34 than in the control (**Figure 1F**). Time-course PEP uptake assays found that ^13^C-PEP was rapidly imported into *E. coli* expressing MBP-SLC25A34 and reached a plateau state within 15 min, whereas no uptake was detected in the control group (**Figure 1G**). To validate the specificity of SLC25A34-mediated PEP transport, we next evaluated ^13^C-PEP import in the presence of excess unlabeled PEP as a competitor (50 mM). We found that ^13^C-PEP uptake was completely blocked by the addition of excess unlabeled PEP (**Figure 1H**). To verify whether the observed PEP uptake activity is specific to SLC25A34, we expressed MBP-SLC25A45 as an additional control. SLC25A45 is a mitochondrial carrier for trimethyl-lysine (TML) that we and others have recently characterized (*40–42*). Under the same experimental conditions as MBP-SLC25A34, the assay found no PEP uptake in MBP-SLC25A45 (**Figure 1I**). To test its substrate specificity, we tested whether SLC25A34 transports oxaloacetate (OAA), using ^13^C_4_-OAA as a tracer. Since conventional LC–MS assays were unable to detect ¹³C₄-OAA due to its instability, we quantified ¹³C₄-malate derivatized with 3-nitrophenylhydrazine (3-NPH), which served as a proxy for ¹³C₄-OAA (*43*). In contrast to PEP, however, we did not find active OAA transport in the system (**Supplementary Figure 1D**).

As a complementary approach, we developed a cell-free proteo-liposome system with purified SLC25A34 protein. In brief, C43 (DE3) bacteria expressing either an MBP-SLC25A34-TwinStrep construct (SLC25A34) or a control vector expressing the endogenous MBP (control) were subjected to affinity purification using StrepTactin resin (**Figure 1J**). Subsequently, the purified eluents (SLC25A34 protein or the corresponding control fraction) were reconstituted into liposomes. Consistent with the bacterial system, we found that ^13^C-PEP transport was significantly higher in the SLC25A34-liposomes preloaded with PEP than in control-liposomes. Background signals obtained from protein-free empty control liposomes (*i.e.,* liposome lipids only) were subtracted, as these signals represent non-specific association with the liposomes (**Supplementary Figure 1E**). The PEP transport was significantly reduced in the presence of an excess amount (50 mM) of unlabeled PEP (**Figure 1K**).

Given the high structural similarity between SLC25A34 and SLC25A35 (see Figure 1A-C), we asked whether SLC25A34 exhibits comparable transport activity and substrate specificity. To this end, we purified SLC25A34 and SLC25A35 proteins and reconstituted them into liposomes. The assay showed active ^13^C-PEP transport at comparable levels in liposomes containing either SLC25A35 or SLC25A34 (**Supplementary Figure 1F**). To determine the substrate specificity of SLC25A34, we tested ^13^C-PEP transport in the presence of excess unlabeled PEP, malate, OAA, citrate, α-ketoisocaproic acid (KIC), and α-ketoisovaleric acid (KIV), according to our recent study of SLC25A35 (*31*). We found that PEP transport was substantially blunted by PEP, whereas OAA, citrate, KIC, and KIV did not compete with PEP transport via SLC25A34 (**Figure 1L** and **Supplementary Figure 1G**). Malate showed modest yet significant inhibition, albeit to a lesser extent than PEP. The results are consistent with the substrate specificity of SLC25A35 (*31*).

Next, we assessed the functional role of the conserved residues identified in our structural analysis of SLC25A34 by generating a recombinant mutant SLC25A34 protein carrying Y125A/R176A/R277A. We then performed PEP transport assays in liposomes containing the wild-type or mutant form of SLC25A34. Unexpectedly, the assay found higher PEP transport in the SLC25A34 mutant (Y125A/R176A/R277A) than in the wild-type form (**Figure 1M**). It is worth noting that we obtained a similar result with SLC25A35, in which mutations at the corresponding substrate-binding residues increased PEP transport activity compared with the wild-type protein (*31*). These results are in alignment with a report on the ADP/ATP carrier showing that mutations weakening the salt-bridge gate network make the gate leaky, thereby increasing the apparent transport rate (*44*). Our proteo-liposome assay was performed under saturating substrate conditions, and the readout reflects V_max_; hence, gate-loosening mutations could lead to higher apparent transport activity. Nonetheless, the mutation analysis supports the structural prediction that Y125, R176, and R277 are functionally important for SLC25A34-mediated PEP transport.

### SLC25A34 expression is induced by fasting and exercise

We next asked how the SLC25A family members are regulated by metabolic stress, such as fasting and exercise, in skeletal muscle. First, data mining of publicly available data (*45*) found that *Slc25a34* was the most upregulated *Slc25a* gene in response to fasting (**Figure 2A**). In contrast, *Slc25a35* mRNA levels were reduced by fasting. We validated this observation using independent samples: *Slc25a34* mRNA levels in the soleus muscle were upregulated following 24 hours of fasting (**Figure 2B**). A similar trend was also observed in the plantaris muscle, although the basal expression levels in the plantaris muscle were lower than those in the soleus muscle (**Supplementary Figure 2A**, **B**). To examine whether such regulation occurs in a cell-autonomous manner, we investigated *Slc25a34* expression in cultured myocytes in which *Slc25a34* mRNA levels were increased during myoblast differentiation (**Figure 2C**). We found that *Slc25a34* mRNA expression was significantly induced when cells were cultured in a starvation medium (HBSS supplemented with 5.5 mM glucose) for 6 hours or longer, compared to cells cultured in a medium containing 25 mM glucose, 2% horse serum, and amino acids, including 4 mM glutamine (**Figure 2D**). Furthermore, data analysis of publicly available RNA-seq data (*46*) showed that *Slc25a34* is the most upregulated *Slc25a* gene in mouse skeletal muscle following exercise (**Figure 2E**) and is also upregulated in BAT upon cold exposure (**Supplementary Figure 2C**). On the other hand, *Slc25a35* expression did not show such an increase in response to exercise and cold exposure. Consistent with the results in mice, transcriptomics data analysis of human skeletal muscle (GSE120862) showed that *SLC25A34* mRNA levels were significantly upregulated 4 hours after one-legged knee extension exercise in untrained males (**Figure 2F**). Together, these results suggest that SLC25A34 is highly expressed in oxidative muscle and its expression is inducible by fasting, exercise, and cold exposure; in contrast, *Slc25a35* expression was distinctly regulated despite both sharing PEP as a specific substrate (**Figure 2G**).

**Figure 2.**
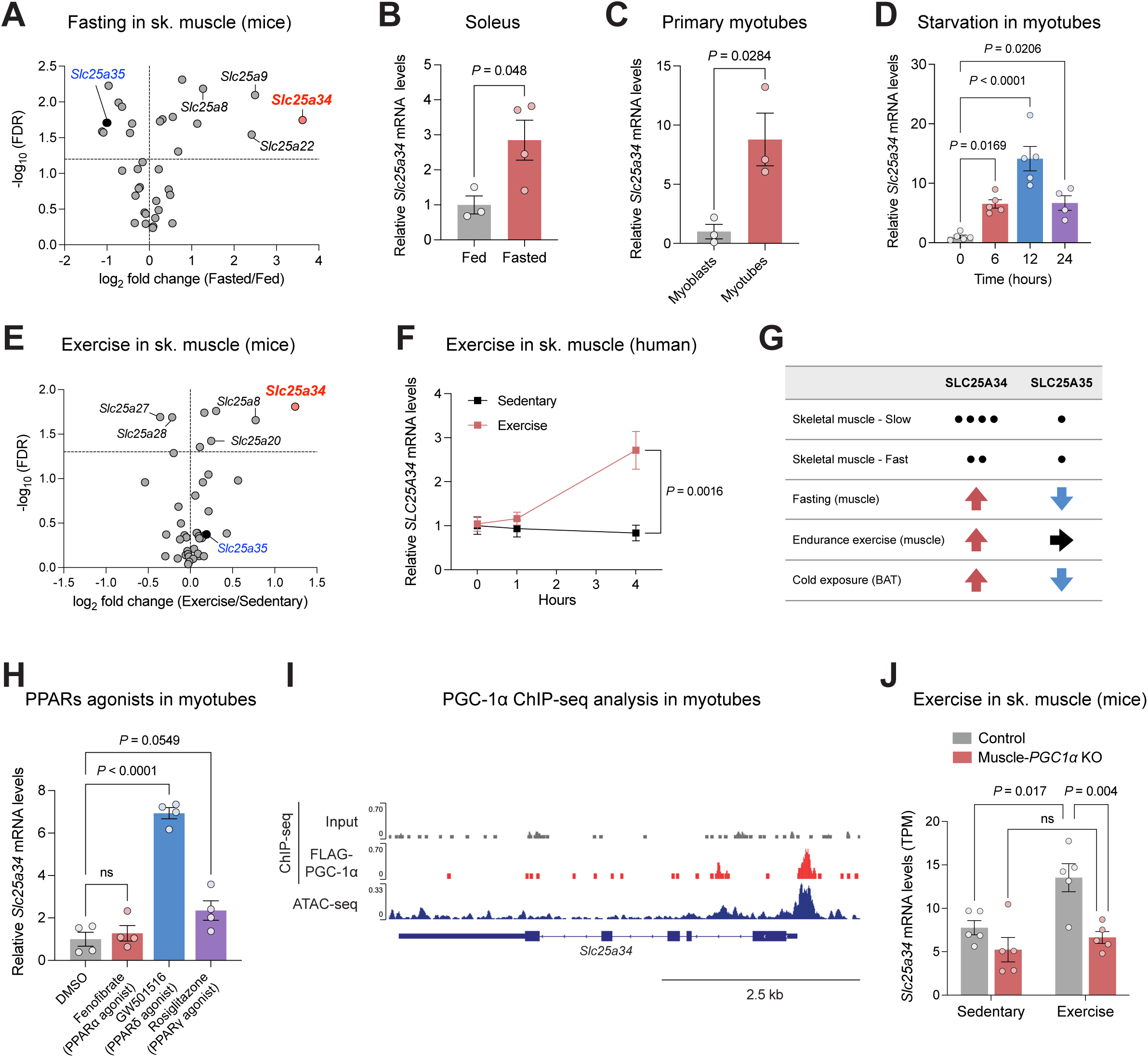
SLC25A34 expression is induced by fasting and exercise. **A.** Transcriptomics of SLC25A genes in the gastrocnemius from fed and 24 hours-fasted mice (GEO: GSE210904). N = 4 per group. **B.** Relative mRNA levels of *Slc25a34* in soleus from wild-type mice after 24 hours of fasting. N = 3 for fed and n = 4 for fasted. **C.** Relative mRNA levels of *Slc25a34* in primary myoblasts and myotubes. N = 3 per group. **D.** Relative mRNA levels of *Slc25a34* in C2C12 myotubes cultured in nutrition-deprived media (HBSS) for indicated times. N = 5 for 0, 6, 12 hours and n = 4 for 24 hours. **E.** Transcriptomics of SLC25A genes in the gastrocnemius of mice after 4 weeks of voluntary running (GEO: GSE123879). N = 5 per group. **F.** Relative mRNA levels of *SLC25A34* in human vastus lateralis from exercised and non-exercised legs after one-legged knee extension (GEO: GSE120862). N = 7 per time point. **G.** Summary of SLC25A34 and SLC25A35 expression and regulation. Dots and arrows indicate relative basal expression and regulatory changes. **H.** Relative mRNA levels of *Slc25a34* in C2C12 myotubes treated with PPAR agonists for 48 hours. N = 4 per group. **I.** The recruitment of PGC-1α to the *Slc25a34* gene locus was visualized from PGC-1α ChIP seq (GEO: GSE51178), and ATAC-seq (GEO: GSE134962). **J.** TPM values of *Slc25a34* in the quadriceps from wild-type and muscle-specific PGC-1α knockout mice after exercise training (GEO: GSE221210). N = 5 per group. Bars represent mean ± s.e.m. *P* values were calculated by unpaired *t*-test with Benjamini-Hochberg FDR correction (A and E), unpaired *t*-test (B and C), one-way ANOVA with Dunnett’s multiple comparisons test (D and H), two-way repeated-measures ANOVA with Tukey’s multiple comparisons test (F) and two-way ANOVA with Tukey’s multiple comparisons test (J).

Next, we examined the regulatory mechanisms of *Slc25a34* expression. One of the key transcriptional regulatory axes mediating the adaptive responses to fasting, exercise, and cold exposure involves peroxisome proliferator-activated receptor-delta (PPARδ) and peroxisome proliferator-activated receptor-gamma coactivator 1-alpha (PGC-1α), as well as AMP-activated protein kinase (AMPK), among other upstream regulators (*47–49*). Notably, treatment with a synthetic agonist for PPARδ (GW501516 at 1 µM), but not agonists for PPARα (fenofibrate, 100 µM) and for PPARγ (rosiglitazone at 10 µM), significantly increased the levels of *Slc25a34* in cultured C2C12 myotubes (**Figure 2H**). GW501516 (a PPARδ agonist) treatment led to a dose- and time-dependent increase in *Slc25a34* in C2C12 myotubes (**Supplementary Figure 2D, E**). Furthermore, data analysis of chromatin immunoprecipitation sequencing (ChIP-seq) of PGC-1α (*50*) showed active recruitment of PGC-1α onto the proximal promoter region as well as the intron of the *Slc25a34* gene locus in C2C12 myotubes (**Figure 2I**). Importantly, transcriptome data analysis of muscle-specific PGC-1α KO mice (*51*) showed that *Slc25a34* mRNA levels were significantly elevated in skeletal muscle of mice following exercise, whereas such induction by exercise was not seen in muscle-specific PGC-1α KO mice (**Figure 2J**). These data indicate that SLC25A34 expression is regulated by the PPARδ/PGC-1α transcriptional cascade in skeletal muscle.

### SLC25A34 is required for amino acid utilization in myocytes

We next examined the metabolic consequence of SLC25A34 loss in myocytes. To this end, we generated primary myoblasts that lacked SLC25A34 (*Slc25a34* KO cells) and control cells (control) by the *Easi*-CRISPR (Efficient additions with ssDNA inserts-CRISPR) system (*52*) (**Figure 3A**). There were no differences in myoblast growth and differentiation between control and *Slc25a34* KO cells in culture (**Supplementary Figure 3A**-**C**). To isolate mitochondria from control and *Slc25a34* KO myotubes rapidly, we expressed the Mito-Tag in these cells (*53*) and analyzed their whole-cell metabolites as well as mitochondrial metabolites by LC-MS (**Figure 3B, Supplementary Figure 3D**). Under standard cultured conditions with 25 mM glucose, 5% horse serum, and amino acids, there was no significant difference in mitochondrial metabolites between control and *Slc25a34* KO myotubes (**Figure 3C, Supplementary Table 1**). Similarly, at the whole-cell level, there was no major change in intracellular PEP contents between the two groups (**Supplementary Figure 3E, Supplementary Table 2**). However, under a starved culture condition with HBSS supplemented with 5.5 mM glucose for 6 hours, we found dynamic differences in mitochondrial metabolites between control and *Slc25a34* KO myotubes. This includes significant reductions in PEP and several TCA intermediates, such as aspartate, malate, fumarate, and αKG; on the other hand, several amino acids (lysine, arginine, glutamine, and threonine) and choline were significantly elevated in *Slc25a34* KO mitochondria (**Figure 3D, Supplementary Table 3**). Such changes were not observed at the whole-cell level (**Supplementary Figure 3F, Supplementary Table 4**). Accordingly, mitochondrial enrichment of PEP in *Slc25a34* KO myotubes was significantly lower than that in control myotubes under this condition (**Figure 3E**). Similarly, mitochondrial enrichment of several metabolites, including malate, fumarate, aspartate, choline, lysine, and arginine, was altered in *Slc25a34* KO cells (**Supplementary Figure 3G**).

**Figure 3.**
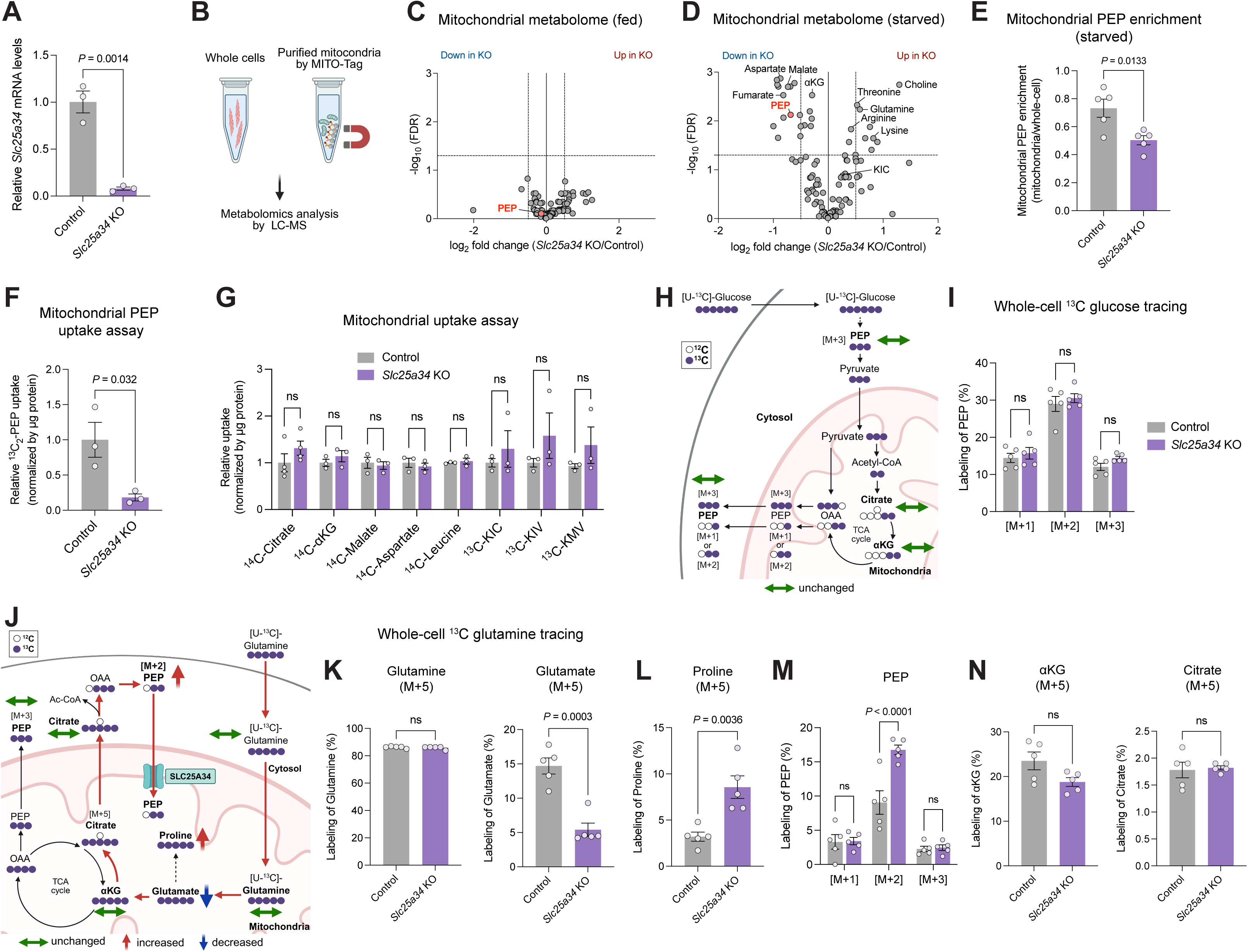
SLC25A34 is required for amino acid utilization in myocytes. **A.** Relative mRNA levels of *Slc25a34* in control and *Slc25a34* KO primary myotubes. N = 3 per group. **B.** Schematic of mitochondrial and whole-cell metabolomics experiments using MITO-Tag. **C.** Mitochondrial metabolomics under a nutrient-replete condition. N = 5 per group. **D.** Mitochondrial metabolomics under a nutrient-depleted condition. N = 5 per group. **E.** Mitochondrial PEP enrichment under a nutrient-depleted condition was calculated as the ratio of mitochondrial to whole-cell PEP intensity. N = 5 per group. **F.** Mitochondrial PEP uptake assay. N = 3 per group. **G.** Mitochondrial uptake of the indicated metabolites. N = 4 for citrate, n = 3 per group for other metabolites. **H.** Schematic of [U-^13^C]-glucose tracing under a fasted condition. Cells were cultured in a nutrient-depleted condition (green arrow, no change). **I.** Labeling of the indicated ^13^C-labeled PEP (%) in (H). n = 5 per group. **J.** Schematic of [U-^13^C]-glutamine tracing. Control and KO primary myotubes were cultured in a nutrient-depleted condition (red arrow, increased; blue arrow, decreased; green arrow, no change). **K.** M+5 glutamine and glutamate labeling (%) in (J). n = 5 per group. **L.** M+5 proline labeling (%) in (J). n = 5 per group. **M.** Labeling of the indicated ^13^C-labeled PEP (%) in (J). n = 5 per group. **N.** M+5 α-ketoglutarate (αKG) and citrate labeling (%) in (J). n = 5 per group. Bars represent mean ± s.e.m. *P* values were calculated by unpaired *t*-test (A, E, F, G, K, L and N), unpaired *t*-test with Benjamini-Hochberg FDR correction (C and D) and two-way ANOVA with Tukey’s multiple comparisons test (I and M).

Since the liposome-based assay suggests that SLC25A34 transports PEP, we next asked whether mitochondrial PEP uptake was altered in *Slc25a34* KO myotubes. To this end, mitochondria isolated from control and *Slc25a34* KO myotubes were incubated with ^13^C_2_-labeled PEP for 5 min, quickly rinsed, and their tracer contents were measured by LC-MS. In alignment with the above results, mitochondrial PEP uptake was significantly reduced in *Slc25a34* KO myotubes compared to that in control cells (**Figure 3F**). To determine the selectivity, we also incubated *Slc25a34* KO and control mitochondria with ^14^C-labeled citrate, αKG, malate, aspartate, leucine, as well as ^13^C-labeled KIC, KIV, and α-keto-β-methylvaleric acid (KMV). However, we did not find any difference in their mitochondrial uptake between the two groups (**Figure 3G**). These results suggest that SLC25A34 loss resulted in the selective impairment of mitochondrial PEP import, whereas changes in mitochondrial enrichment of malate, aspartate, αKG, and other metabolites observed in *Slc25a34* KO myotubes were consequences of *Slc25a34* deletion, rather than reduced mitochondrial import *per se*.

Considering the diverse biological roles of PEP, we next examined the extent to which SLC25A34 loss affects the utilization of glucose, pyruvate, and amino acids by performing tracer studies. First, [U-^13^C]-glucose tracing in differentiated myotubes found no significant difference in M+3, M+2, and M+1 PEP contents between control and *Slc25a34* KO cells (**Figure 3H, I**). Similarly, there was no difference in ^13^C-labeled contents of citrate, αKG, succinate, aspartate, and glutamate between the two groups (**Supplementary Figure 4A**). When we used [U-^13^C]-pyruvate as a tracer, we did not find significant differences in M+3, M+2, and M+1 PEP contents, as well as ^13^C-labeled TCA cycle intermediates, between control and *Slc25a34* KO cells (**Supplementary Figure 4B-D**). These results suggest that SLC25A34 loss does not affect glycolysis or the cellular utilization of glucose and pyruvate in myotubes. This finding is consistent with the cellular respiration data showing that the oxygen consumption rate (OCR) of *Slc25a34* KO myotubes was comparable to that of control cells in cultured media containing high glucose (**Supplementary Figure 4E**).

On the other hand, we found that amino acid utilization was significantly altered in *Slc25a34* KO cells, as summarized in **Figure 3J**. When [U-^13^C]-glutamine was used as a tracer, M+5 glutamate levels were significantly lower in *Slc25a34* KO cells than in control cells, even though M+5 glutamine contents were comparable between the two groups (**Figure 3K**). Data for M+1 to M+4 glutamine and glutamate are shown in **Supplementary Figure 4F, G**. Instead, we found that *Slc25a34* KO cells contained significantly higher levels of M+5 proline than control cells (**Figure 3L**). No change was seen in M+1 to M+4 proline levels between KO and control cells (**Supplementary Figure 4H**). These results suggest that glutamine-driven anaplerosis was impaired, whereas the conversion to proline was enhanced in *Slc25a34* KO cells. Notably, M+2 PEP levels, i.e., PEP regeneration from OAA, but not M+1 and M+3 PEP levels, were higher in *Slc25a34* KO cells than in control cells (**Figure 3M**). Since there was no difference in M+5 αKG and citrate levels between the two groups (**Figure 3N**), the data suggest that glutamine-derived PEP (M+2) is accumulated in *Slc25a34* KO cells due to impaired PEP import into the mitochondria (pathways highlighted in red lines in Figure 3J).

### SLC25A34 loss results in muscle hypertrophy and myopathy

The above results suggest that SLC25A34 is essential for amino acid utilization in skeletal muscle under conditions of nutritional deprivation. Of note, SLC25A34 expression was downregulated in muscle diseases. For example, Duchenne muscular dystrophy (DMD), caused by loss-of-function mutations in the dystrophin gene, is characterized by chronic muscle degeneration and regeneration, leading to alterations in muscle amino acid metabolism (*54, 55*). In a mouse model of DMD (GSE162455), *Slc25a34* mRNA levels in the soleus muscle were lower than in control mice (**Figure 4A**). Furthermore, *SLC25A34* mRNA levels in the vastus lateralis muscle of DMD patients were significantly lower compared to healthy subjects (*56*) (**Figure 4B**). Friedreich’s ataxia (FRDA) is an inherited mitochondrial disease caused by frataxin deficiency, resulting in metabolic skeletal muscle impairments, including downregulation of mitochondrial enzymes involved in amino acid catabolism (*57, 58*). Transcriptomic analysis of Friedreich’s ataxia subjects (*59*) found lower levels of *SLC25A34* mRNA in the gastrocnemius muscle compared to those in healthy control subjects (**Figure 4C**). These results motivated us to investigate how the loss of SLC25A34 in skeletal muscle affects skeletal muscle function *in vivo*.

**Figure 4.**
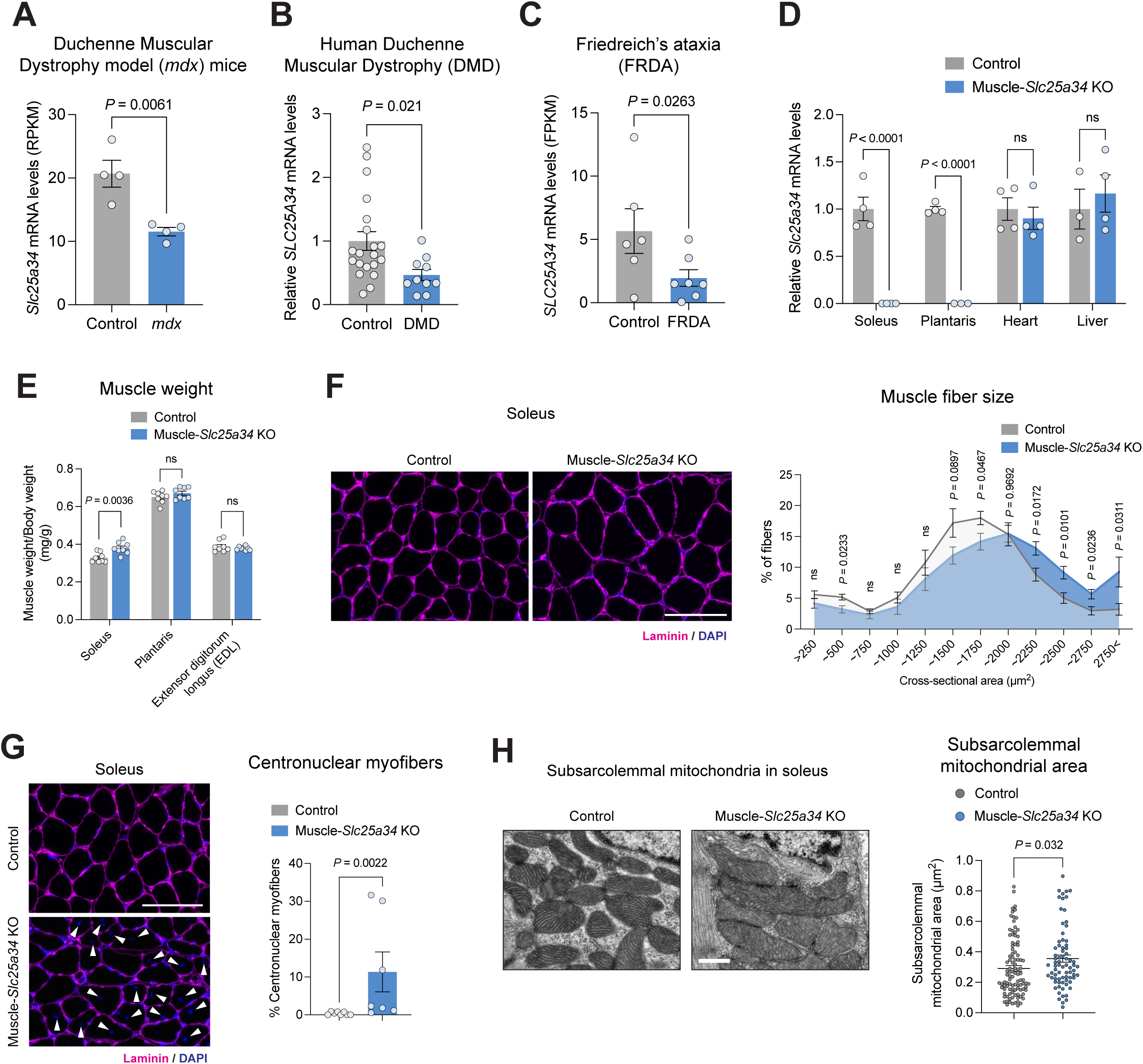
SLC25A34 loss results in hypertrophy and myopathy in slow-twitch muscle. **A.** RPKM values for *Slc25a34* in soleus from *mdx* mice (GEO: GSE162455). N = 4 per group. **B.** Relative *SLC25A34* mRNA levels in human vastus lateralis from healthy and Duchenne Muscular Dystrophy (DMD) subjects (GEO: GSE3307). N = 20 for healthy and n = 10 for DMD subjects. **C.** *SLC25A34* transcript levels in human gastrocnemius from subjects with Friedreich’s ataxia (FRDA) (GEO: GSE226646). N = 6 for control and n = 7 for FRDA subjects. **D.** Relative mRNA levels of *Slc25a34* in the indicated tissues in male control and KO mice. N = 3-4 for control and KO. **E.** Muscle weight of the soleus, plantaris, and extensor digitorum longus (EDL) in male control and KO mice at 16 weeks old. Muscle weight (mg) was normalized to the body weight (g). n = 8 per group. **F.** Representative laminin and DAPI-stained soleus sections from male mice. Right: Distribution of the cross-sectional area. Scale bar, 100 µm. n = 7 for control and n = 6 for KO. **G.** Representative laminin and DAPI-stained soleus sections from male mice. Central nuclei are indicated by arrowheads. Right: Quantification of centrally nucleated myofibers per soleus cross-section. Scale bar, 100 µm. n = 8 for control and n = 7 for KO mice. **H.** Representative electron microscopy images of subsarcolemmal mitochondria in soleus of male mice. Right: Quantification of subsarcolemmal mitochondrial area. Scale bar, 0.5 µm. n = 101 for control and 75 for KO collected from n = 2 mice per group. Bars represent mean ± s.e.m. *P* values were calculated by unpaired *t*-test (A, B, D, E, F and H), Wald test with Benjamini-Hochberg FDR correction (C) and Mann-Whitney test (G).

To this end, we developed *Slc25a34*^flox/flox^ mice and subsequently crossed them with α-skeletal actin (HSA)-Cre mice (HSA-Cre x *Slc25a34*^flox/flox^, herein muscle-*Slc25a34* KO mice). We validated that HSA-Cre effectively deleted *Slc25a34* in the soleus and plantaris muscles of muscle-*Slc25a34* KO mice, but not in the heart and liver (**Figure 4D**). We found no difference in body weight between muscle-*Slc25a34* KO mice and littermate control mice (*Slc25a34*^flox/flox^) when mice were fed a regular diet at room temperature (**Supplementary Figure 5A**). However, the soleus muscle of muscle-*Slc25a34* KO mice exhibited modest but significantly greater tissue mass than that of control mice (**Figure 4E**). This phenotype was consistently observed both in male and female mice (**Supplementary Figure 5B**). Note that the tissue weight of the soleus muscle was greater in muscle-*Slc25a34* KO mice than in control mice without normalization to body weight (**Supplementary Figure 5C**). The difference was attributed to myofiber hypertrophy, as the soleus muscle of muscle-*Slc25a34* KO mice contained larger myofibers than those of control mice (**Figure 4F**). On the other hand, we did not find changes in the composition of muscle fiber types (Type I, IIa, IId/x) between muscle-*Slc25a34* KO mice and littermate control mice (**Supplementary Figure 5D**). In alignment with the results in cellular respiration (shown in Supplementary Figure 4E), there was no significant difference in muscle tissue oxygen consumption rate (*J*O_2_) between the two groups when pyruvate, malate, and succinate were provided (**Supplementary Figure 5E**). Additionally, muscle-*Slc25a34* KO mice showed endurance capacity and grip strength similar to control mice (**Supplementary Figure 5F, G**).

However, we found that the muscle-*Slc25a34* KO mice exhibited two notable abnormalities in the soleus muscle. First, there was a marked increase in centronuclear myofibers, defined by the presence of centrally located myonuclei, rather than the normal peripheral localization beneath the sarcolemma (arrowheads in the left panel of **Figure 4G**). In contrast to healthy adult muscle fibers in control mice, where nuclei resided at the periphery, muscle-*Slc25a34* KO mice contained centronuclear fibers, which are recognized as a morphological hallmark of active muscle regeneration, also reflecting cycles of degeneration or chronic myofiber stress (right panel in **Figure 4G**; quantification is shown on the right graph). Second, the muscle-*Slc25a34* KO mice displayed enlarged subsarcolemmal mitochondria in the soleus (**Figure 4H**). It is worth noting that this mitochondrial enlargement was uniquely found in the subsarcolemmal compartment, but was not observed in the intermyofibrillar mitochondrial network (**Supplementary Figure 5H**). This compartment-selective phenotype aligns with prior studies showing that subsarcolemmal mitochondria exhibit distinct metabolic sensitivities to amino acid-related stimuli, such as increased ADP-stimulated respiration and ATP production following acute amino acid infusion in humans (*60*), and augmented glutamate-driven respiration after endurance exercise training (*61*). These results suggest that loss of SLC25A34 resulted in abnormalities in the ultrastructure of subsarcolemmal mitochondria in the soleus muscle, contributing to localized metabolic stress and regenerative responses.

### Altered protein turnover in the soleus muscle of *Slc25a34* KO mice

To determine the molecular changes caused by SLC25A34 loss in the soleus muscle, we next performed RNA-sequencing in muscle-*Slc25a34* KO and littermate control mice. The transcriptome analysis identified 680 genes that were significantly altered between the control and muscle-*Slc25a34* KO groups (131 genes were upregulated, 549 genes were downregulated). Among these changes, the MSigDB hallmark gene sets analysis found that the myogenesis/regenerative gene signature (e.g., *Mylpf, Myod1*, *Myh8,* and *Tnnt2*) and IL-6/JAK/STAT3 signaling (e.g., *Socs3*) were significantly upregulated in muscle-*Slc25a34* KO mice relative to control mice (**Figure 5A, B**). On the other hand, several genes known for the regulation of PI3K/AKT/mTOR signaling (e.g., *Pten* and *Tsc1*, negative regulators of mTOR signaling), were significantly downregulated in muscle-*Slc25a34* KO mice relative to control mice. These transcriptional changes appeared to be independent of general mitochondrial dysfunction *per se*, as many, if not all, of the mitochondrial TCA cycle genes and OXPHOS proteins were expressed at equivalent levels in both groups (**Supplementary Figure 6A, B**).

**Figure 5.**
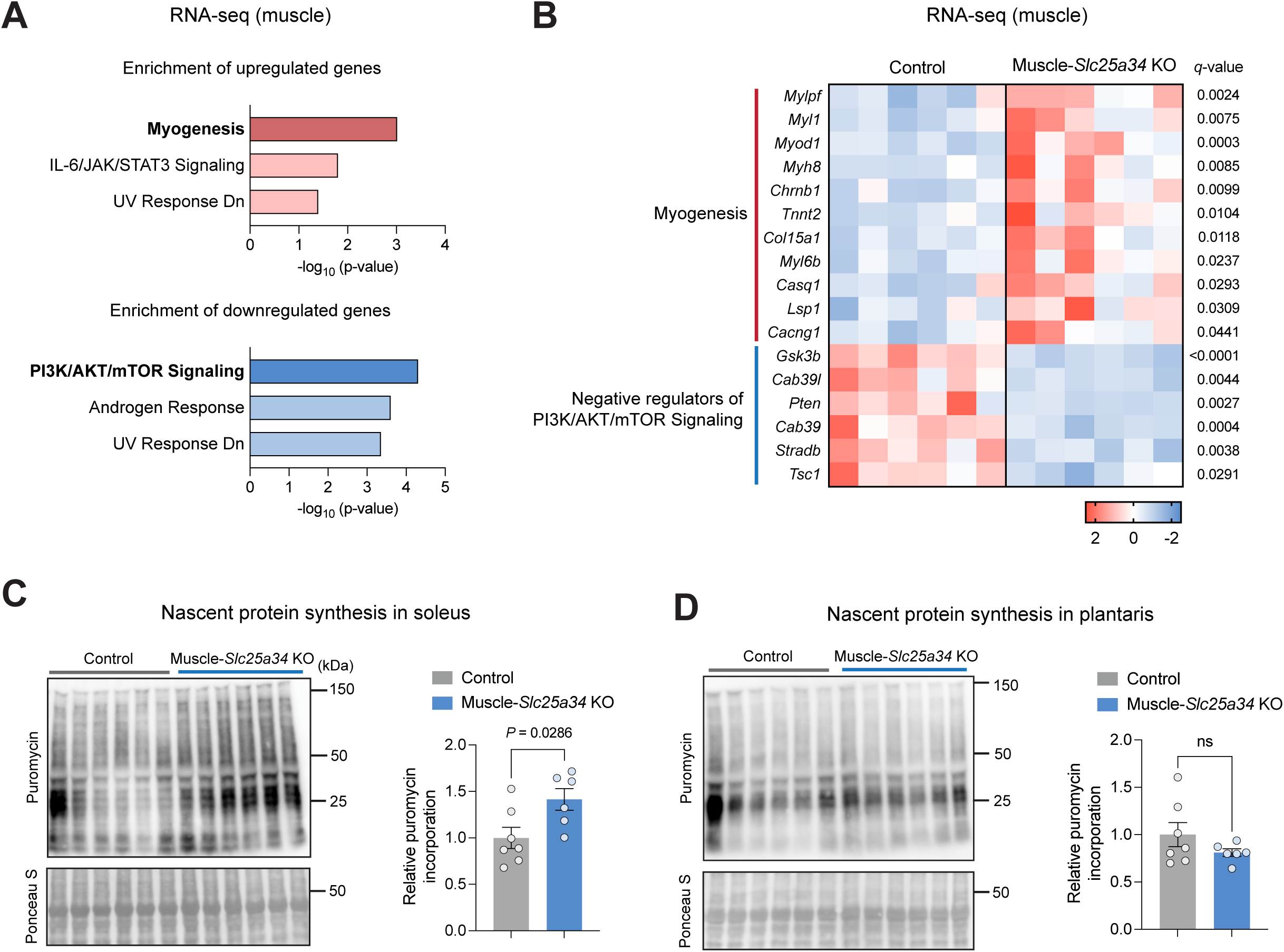
Altered protein turnover in the soleus muscle of *Slc25a34* KO mice. **A.** MsigDB hallmark gene set enrichment analysis (Enrichr) was performed using genes that were significantly (*q* < 0.01) upregulated or downregulated in the *Slc25a34* KO soleus muscle compared to the control in RNA-seq analysis. **B.** Relative expression levels of genes related to myogenesis and PI3K/AKT/mTOR signaling in the soleus muscle of control and muscle-*Slc25a34* KO mice. Data are represented as *Z*-score heatmaps for the indicated genes in each sample, based on RNA-seq analysis. N = 6 per group. Red indicates elevated levels, and blue indicates decreased levels. **C.** Nascent protein synthesis in the soleus muscle of female control and muscle-*Slc25a34* KO mice, assessed by the SUnSET assay. Immunoblotting was performed to detect the puromycin. The intensity of puromycin was normalized to the total protein level (Ponceau S staining). The molecular weights (kDa) are indicated on the right. N = 7 for control and 6 for muscle-*Slc25a34* KO. **D.** Nascent protein synthesis in the plantaris muscle of female control and muscle-*Slc25a34* KO mice, assessed by the SUnSET assay. Immunoblotting was performed to detect the puromycin. The intensity of puromycin was normalized to the total protein level (Ponceau S staining). The molecular weights (kDa) are indicated on the right. N = 7 for control and 6 for muscle-*Slc25a34* KO. Bars represent mean ± s.e.m. *P* values were calculated by Fisher’s exact test (A), unpaired *t*-test (C and D), and *q*-values were calculated by Wald test with Benjamini-Hochberg FDR correction (B).

The PI3K/AKT/mTOR signaling pathway plays a central role in integrating nutrient and hormonal cues to drive anabolic protein synthesis (*62*). Down-regulation of negative regulators of mTOR signaling suggested that elevated muscle protein synthesis was accompanied by active muscle regeneration and tissue remodeling in muscle-*Slc25a34* KO mice. To test this possibility, we performed the SUnSET assay, a puromycin-based method to assess nascent protein synthesis *in vivo* (*63*). Control and muscle-*Slc25a34* KO mice received an intraperitoneal injection of puromycin (0.04 µmol/g body weight), and the soleus and plantaris muscles were harvested exactly 30 min after puromycin injection. We found that puromycin incorporation into nascent polypeptides was significantly elevated in the soleus muscle of muscle-*Slc25a34* KO mice relative to control mice (**Figure 5C**). This change was selective to the soleus muscle rather than a global change in protein synthesis because there was no difference in puromycin incorporation into nascent proteins in the plantaris muscle between the two groups (**Figure 5D**). These results suggest that muscle hypertrophy in the soleus of muscle-*Slc25a34* KO mice is due to an elevated regeneration program and protein synthesis.

### SLC25A34 loss leads to increased protein synthesis via the mTORC1 pathway

We asked to what extent the muscle hypertrophy observed in muscle-*Slc25a34* KO mice was due to cell-autonomous changes in myofibers. Under a nutrient-replete culture condition (25 mM glucose, 2% horse serum, and amino acids, including 4 mM glutamine), there was no statistically significant difference in myotube diameter between control and *Slc25a34* KO myotubes (**Figure 6A**, left panels). When cells were cultured in a nutrient-deprived medium with HBSS supplemented with 5.5 mM glucose for 6 hours, control myotubes became noticeably thinner, and their diameter was significantly reduced relative to the fed condition, consistent with increased protein degradation during fasting (**Figure 6A**, right panels). *Slc25a34* KO myotubes were significantly resistant to this starvation-induced reduction in myotube diameter, indicating a defect in proteostasis under a nutrient-deprived condition (**Figure 6B**). Since SLC25A35 shares the same substrate as SLC25A34, we next asked whether SLC25A35 also plays a similar biological role in starvation-induced myotube atrophy and amino acid metabolism. Of note, *Slc25a35* mRNA levels were much lower than those of *Slc25a34* in skeletal muscle (see **Supplementary Figure 1A**) and were not induced by fasting (see **Figure 2A**). We isolated myotubes from *Slc25a35* KO mice and cultured them under a nutrient-deprived medium (HBSS supplemented with 5.5 mM glucose) for 6 hours, under the same protocol for *Slc25a34* KO myotubes (**Supplementary Figure 7A**). In contrast to *Slc25a34* KO myotubes, *Slc25a35* deficiency did not affect atrophy in response to starvation (**Figure 6C**).

**Figure 6.**
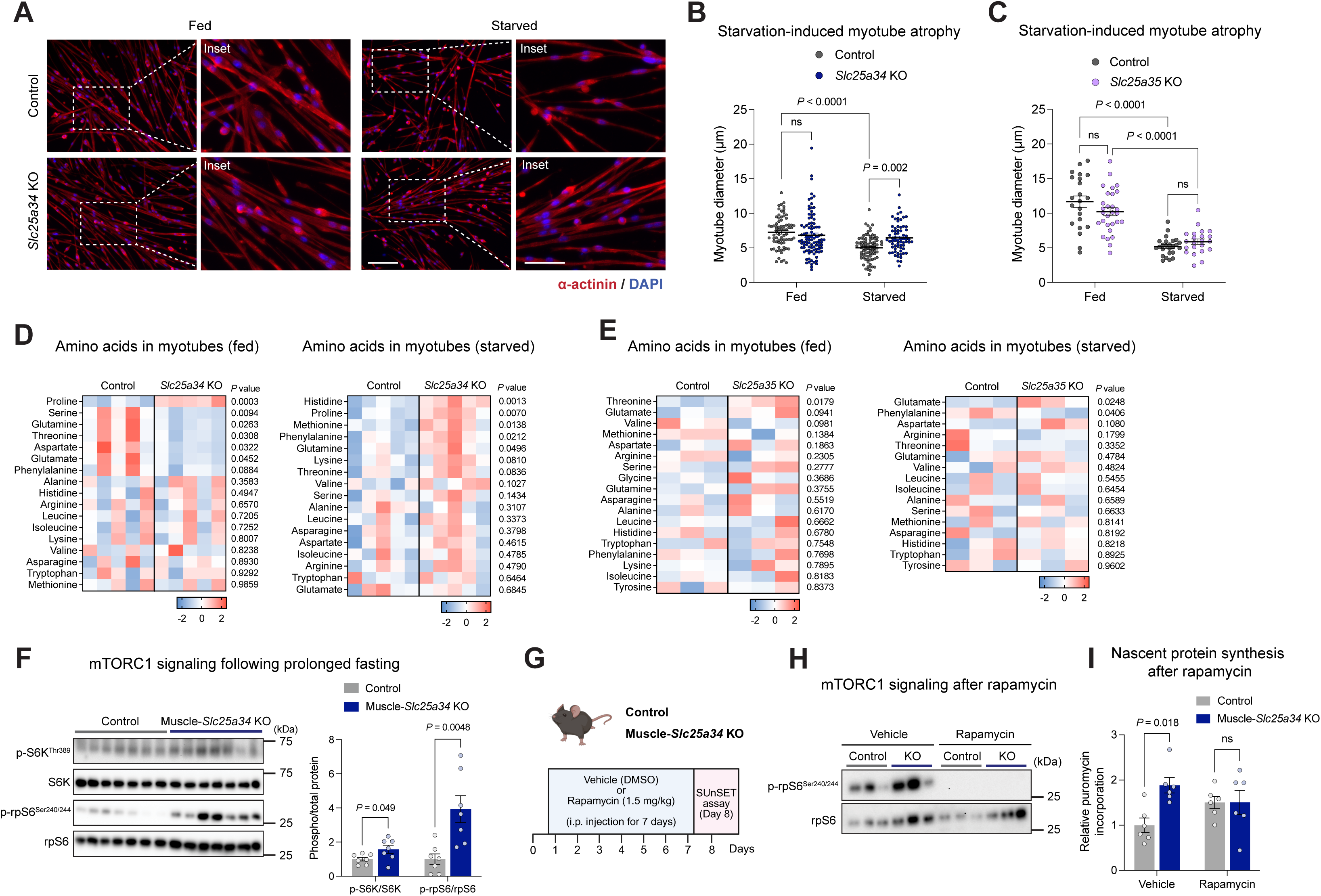
SLC25A34 loss leads to increased protein synthesis via the mTORC1 pathway. **A.** Representative α-actinin and DAPI-stained myotubes. Left: Cells were cultured in a nutrient-replete medium. Right: Cells were cultured in a nutrient-deprived medium. Scale bar, 100 µm. Insets show higher-magnification views of the boxed regions. Scale bar, 50 µm. **B.** Starvation-induced myotube atrophy in control and *Slc25a34* KO primary myotubes. N = 74 for control-fed, n = 83 for control-starved, n = 95 for KO-fed, n = 67 for KO-starved. **C.** Starvation-induced myotube atrophy in control and *Slc25a35* KO primary myotubes. N = 22 for control-fed, n = 25 for control-starved, n = 29 for KO-fed, n = 22 for KO-starved. **D.** Relative amino acid levels in control and *Slc25a34* KO myotubes in a nutrient-replete medium, and in a nutrient-deprived medium. N = 5 per group. **E.** Relative amino acid levels in control and *Slc25a35* KO myotubes in a nutrient-replete medium, and in a nutrient-deprived medium. N = 3 per group. **F.** mTORC1 signaling after 24 hours of fasting. Immunoblotting was performed to detect p-S6K^Thr389^, total S6K, p-rpS6^Ser240/244^ and total rpS6. N = 7 per group. **G.** Schematic of SUnSET assay following the inhibition of mTORC1 signaling. **H.** mTORC1 signaling in the soleus muscle of male control and muscle-*Slc25a34* KO mice following rapamycin treatment. Immunoblotting was performed to detect p-rpS6^Ser240/244^ and total rpS6. **I.** Nascent protein synthesis in the soleus muscles of male control and muscle-*Slc25a34* KO mice following rapamycin treatment. n = 6 per group. Bars represent mean ± s.e.m. *P* values were calculated by two-way ANOVA with Tukey’s multiple comparisons test (B, C, I) and unpaired *t*-test (D, E, F).

Consistent with these morphological changes, we found differences in intracellular amino acid levels between control and *Slc25a34* KO myotubes in a nutritional state-dependent manner. Metabolomics found elevated cellular proline levels in *Slc25a34* KO cells compared to control cells (**Figure 6D**, left panel, **Supplementary Table 5**). In addition, several amino acids, including serine, glutamine, threonine, aspartate, and glutamate, were lower in *Slc25a34* KO cells than in control cells. Under a nutrient-deprived condition, several amino acid levels were significantly elevated in *Slc25a34* KO cells compared to control cells, including proline and glutamine (**Figure 6D**, right panel, **Supplementary Table 6**). In contrast, intracellular amino acid levels were largely unchanged in *Slc25a35* KO myotubes compared with control myotubes under both fed and starved conditions (**Figure 6E, Supplementary Tables 7 and 8**). Together, these results indicate that SLC25A34 plays a distinct functional role in maintaining muscle integrity during nutritional deprivation, whereas SLC25A35 is dispensable, despite the two sharing PEP as a substrate.

Since these amino acids are known activators of the mTOR complex (*64*), and our RNA-seq data showed that several genes known for negatively regulating mTOR signaling, including *Pten* and *Tsc1*, were down-regulated in muscle-*Slc25a34* KO mice (see Figure 5B), these data collectively led to the hypothesis that mTORC1 signaling is activated in the absence of SLC25A34 even under a nutrition-deprived state. To test this, we harvested the soleus muscle of control and muscle-*Slc25a34* KO mice after 24 hours of fasting. Immunoblotting showed that phosphorylated S6K^Thr389^ and phosphorylated rpS6^Ser240/244^, both of which are established markers of mTORC1-S6K signaling, were significantly elevated in the soleus muscle of muscle-*Slc25a34* KO mice compared with controls (**Figure 6F**). Because AMPK serves as a key sensor of cellular energy status and an upstream negative regulator of mTORC1 signaling (*49, 65*), we next examined AMPK signaling in the soleus muscle of muscle-*Slc25a34* KO mice. However, we found that phosphorylated AMPKα^Thr172^ levels were comparable between control and muscle-*Slc25a34* KO soleus muscles after 24 hours of fasting (**Supplementary Figure 7B**).

Lastly, we examined the functional contribution of elevated mTOR signaling to the increased protein synthesis observed in muscle-*Slc25a34* KO mice. To this end, we treated muscle-*Slc25a34* KO mice and littermate control mice with rapamycin (1.5 mg/kg) for 7 days via *i.p.* injection (**Figure 6G**). After the treatment, we examined mTOR signaling as well as nascent protein synthesis in the soleus muscle. As anticipated, rapamycin treatment effectively reduced the levels of phosphorylated rpS6^Ser240/244^ in both control and muscle-*Slc25a34* KO mice (**Figure 6H**). Consistent with the above result (see Figure 5C), the SUnSET assay showed that vehicle-treated muscle-*Slc25a34* KO mice showed elevated protein synthesis in the soleus compared to control mice that were treated with vehicle. However, rapamycin treatment for 7 days blunted the difference in protein synthesis between muscle-*Slc25a34* KO mice and control mice (**Figure 6I** and **Supplementary Figure 7C**). Rapamycin-treated muscle-*Slc25a34* KO mice and control mice exhibited comparable endurance capacity and grip strength (**Supplementary Figure 7D, E**). These results suggest that elevated amino acids and subsequent activation of mTORC1 signaling mediate active protein synthesis observed in the soleus muscle of muscle-*Slc25a34* KO mice.

## DISCUSSION

Skeletal muscle is the largest protein reservoir in the body and plays a central role in metabolic adaptation during fasting. Under the conditions of nutrient deprivation, muscle increases proteolysis to supply amino acids for hepatic gluconeogenesis and for use by other metabolic organs, while simultaneously oxidizing amino acids via anaplerotic pathways to sustain mitochondrial ATP synthesis. These adaptive responses are accompanied by suppression of mTORC1 signaling, reduced protein synthesis, and enhanced protein breakdown (*7, 8*). In this study, we identify SLC25A34 as a fasting- and exercise-inducible mitochondrial carrier that is required for maintaining amino acid catabolism in skeletal muscle (**Figure 7**). SLC25A34 expression is induced by the PPARδ/PGC-1α transcriptional axis, linking its regulation to established programs of oxidative and endurance adaptation. Consistent with our recent identification of SLC25A35 as a mitochondrial PEP carrier (*31*), we found that SLC25A34 also mediates PEP transport into the mitochondrial matrix. SLC25A34-dependent PEP import in muscle supports glutamine-driven anaplerosis and preserves the balance between TCA cycle anaplerosis and cataplerosis during fasting. Loss of SLC25A34 resulted in blunted amino acid catabolism, leading to amino acid accumulation and elevated mTORC1 signaling under the conditions of nutritional deprivation. Importantly, muscle-specific deletion of *Slc25a34* resulted in elevated protein synthesis, hypertrophy, and myopathy in the soleus muscle, whereas the increased protein synthesis can be reversed by mTORC1 inhibition. Together, the present study identifies SLC25A34 as a mitochondrial metabolic hub that preserves amino acid utilization and proteostasis in skeletal muscle under conditions of nutrient deprivation.

**Figure 7.**
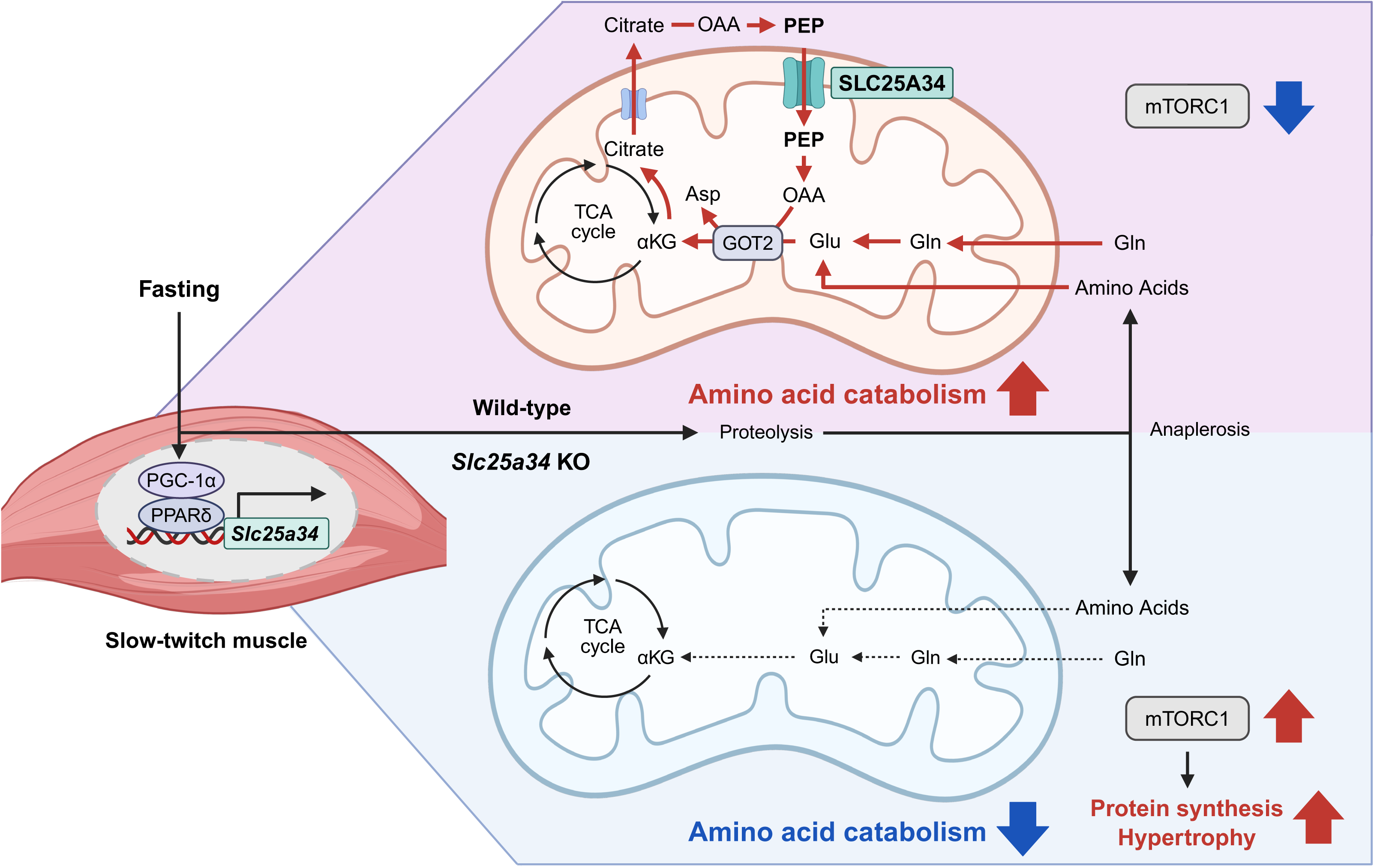
A model of SLC25A34-mediated PEP transport across the IMM regulates amino acid catabolism and maintains metabolic resilience in skeletal muscle during fasting. Under conditions of limited glucose availability, such as prolonged fasting, skeletal muscle relies on amino acids to sustain anaplerotic flux, accompanied by suppression of nutrient-sensing mTORC1 signaling and reduced protein synthesis. SLC25A34 expression is strongly induced by fasting through the PPARδ/PGC-1α transcriptional pathway. SLC25A34-dependent PEP import into mitochondria is essential for the balance between TCA cycle anaplerosis and cataplerosis during fasting; loss of SLC25A34 reduces mitochondrial PEP availability and impairs amino acid catabolism (indicated by red lines). As a consequence, SLC25A34 deficiency leads to amino acid accumulation, inappropriate activation of mTORC1 signaling, and enhanced protein synthesis, leading to hypertrophy and myopathic remodeling in the soleus muscle. Pharmacological inhibition of mTORC1 fully reverses enhanced protein synthesis, demonstrating that aberrant mTORC1 activation is the key driver of the anabolic remodeling in muscle lacking SLC25A34.

Broadly, our findings underscore the importance of coordinated anaplerotic and cataplerotic balance in skeletal muscle metabolism. During prolonged fasting, precise regulation of these pathways via SLC25A34 is necessary to maintain the TCA cycle intermediate pool by utilizing amino acids as alternative substrates. Disruption of this balance leads to the depletion or inappropriate accumulation of TCA intermediates and their precursors, with deleterious consequences for muscle integrity. Consistent with this principle, mutations in enzymes or transporters that regulate anaplerotic or cataplerotic flux are associated with muscle pathology and human disease. For example, SUCLA2, which encodes the β-subunit of ADP-forming succinyl-CoA synthetase, is required for efficient utilization of anaplerotic carbon entering the TCA cycle via amino acid-derived propionyl-CoA. Mutations in SUCLA2, which is highly expressed in skeletal muscle and brain, cause Leigh-like encephalomyopathy accompanied by elevated methylmalonic acid levels (*66, 67*). Loss-of-function mutations in SLC25A1, a mitochondrial citrate carrier that mediates cataplerotic export of citrate from the TCA cycle (*68*), is associated with congenital myasthenic syndrome characterized by fatigable proximal muscle weakness, enlarged mitochondria, and mitochondrial abnormalities (*69, 70*). Additionally, biallelic mutations in SLC25A10, which impair anaplerotic replenishment of malate and succinate, lead to severe epileptic encephalopathy with mitochondrial dysfunction in skeletal muscle (*71*).

We note two limitations in the present study. First, it remains unknown which metabolites mediate mTORC1 activation in muscle-*Slc25a34* KO muscle. Several amino acids, including Met and Gln that activate mTORC1 signaling (*72, 73*), were elevated in *Slc25a34* KO myotubes, whereas canonical mTORC1-activating amino acids (Leu, Arg) were unchanged. It is conceivable that Met via SAMTOR signaling mediates mTORC1 activation in *Slc25a34*-deficient muscles. The second is that the liposome-based assays could not determine the precise affinity of SLC25A34 relative to SLC25A35 due to the methodology we used, which required a millimolar concentration of ^13^C-PEP, substantially higher than physiologically relevant PEP concentrations (50-60 µM in HeLa cells). Additionally, the use of ultracentrifugation for separation limits the temporal resolution of PEP transport kinetics.

Nonetheless, our study provides new insights into a broader principle of mitochondrial metabolite transport: several members of the SLC25A carrier family share overlapping substrate specificities, i.e., “redundant” carriers, yet fulfill distinct biological roles that arise from their tissue-specific expression patterns, regulatory mechanisms, and the unique metabolic demands in a given cell/tissue type. Despite SLC25A34 and SLC25A35 sharing PEP as a substrate, SLC25A34 performs a specialized, non-redundant function in skeletal muscle during nutritional stress, as SLC25A34, but not SLC25A35, is highly inducible by fasting, exercise, and cold exposure. Functionally, SLC25A35 is required for glycerolipid synthesis in lipogenic cells (*31*), whereas SLC25A34 is required for amino acid catabolism in the muscle in response to fasting. Similarly, the aspartate/glutamate carriers AGC1 (SLC25A12) and AGC2 (SLC25A13), which mediate exchange of aspartate for glutamate and protons across the IMM as part of the malate–aspartate shuttle (*25*), have distinct biological roles: AGC1 is highly expressed in skeletal and cardiac muscle and is essential for neuronal myelination (*74*), while AGC2 is ubiquitously expressed and regulates hepatic urea cycle function (*75*). Accordingly, mutations in SLC25A12 cause developmental and epileptic encephalopathies (*76*), whereas mutations in SLC25A13 result in citrullinemia type II and neonatal intrahepatic cholestasis (*75*). A similar distinction exists between ANT isoforms: ANT2 (SLC25A5) is ubiquitously expressed and essential for cellular viability, with deficiency causing embryonic lethality in mice (*77*), whereas ANT1 (SLC25A4) is enriched in energetically demanding tissues such as skeletal muscle and heart, where its loss leads to mitochondrial myopathy and cardiomyopathy in mice (*78*) and humans (*79*). Collectively, the present study reinforces the notion that mitochondrial carriers are not interchangeable conduits but are tailored to meet cell- and tissue-specific metabolic demands. Understanding such functional specialization will be crucial for deciphering mitochondrial contributions to metabolic adaptation, muscle disease, and systemic energy homeostasis, extending beyond ATP production.

## MATERIALS AND METHODS

### Animal study

All the animal experiments in this study were performed in compliance with protocols approved by the Institutional Animal Care and Use Committee (IACUC, Protocol# 028-2022-25) at Beth Israel Deaconess Medical Center. Mice were kept under a 12 hr:12 hr light-dark cycle at ambient temperature (22-23°C) and had free access to food and water. For the fasting experiments, the food was removed from cages of fasted animals for 24 hours. *Slc25a34*^flox/flox^ mice on a C57BL/6J background were generated by the *Easi*-CRISPR method at the BIDMC Transgenic Core Facility (*52*). Skeletal muscle-specific *Slc25a34* knockout mice were obtained by crossing the *Slc25a34*^flox/flox^ mice with human α-skeletal actin (HSA)-Cre mice (B6.Cg-Tg (ACTA1-cre)79Jme/J, 006149 The Jackson Laboratory). A list of the ssDNA, gRNAs and genotyping primers sequences is provided in **Supplementary Table 9**.

### Cells

C2C12 myoblasts were cultured with 20% FBS in DMEM at subconfluence. After myoblasts reached confluence, the medium was switched to 2% horse serum (heat-inactivated HS, Gibco) in DMEM for the differentiation. After 5-7 days of differentiation, myotubes were treated with PPARδ agonist (GW501516, SML1491, Sigma), PPARα agonist (fenofibrate, F6020, Sigma) or PPARγ agonist (rosiglitazone, 71740, Cayman Chemical) at indicated dose and time. For experiments that required starvation, differentiated myotubes were washed twice with PBS and incubated in Hanks’ Balanced Salt Solution (HBSS) medium (H8264, Sigma).

To isolate primary myoblasts, hindlimb muscles of *Slc25a34*^flox/flox^ mice or *Slc25a35*^flox/flox^ mice (1 week-old) were isolated and digested with digestion buffer containing Collagenase, type 2 (0.2%, Worthington Biochemical Corporation), Dispase II (2.4U/mL, Millipore), and 2.5 mM CaCl_2_ at 37°C for 45 min. The digested cell mixture was passed through a 100 µm and a 40 µm filter to remove cell debris. Cells were collected by centrifugation for 5 min at 350 x g and suspended in Ham’s F-10 nutrient mixture supplemented with 20% FBS and 2.5 ng/mL human recombinant bFGF (Stem Cell Technology). After 4 hours of incubation, the medium containing cells was transferred to collagen-coated plates and cultured overnight and then transferred to new collagen-coated plates again to purify primary myoblasts (pre-plating). At 60-80% confluence, muscle differentiation was induced by replacing the medium with 5% HS in DMEM. After 2 days of incubation, myotubes were infected with Ad-CMV-eGFP (1060, Vector Biolabs) or Ad-CMV-iCre (1045, Vector Biolabs) overnight and analyzed 2 days after the infection. For starvation experiments, cells were washed twice with PBS and cultured with HBSS (Sigma) for 6 hours. The fusion index was determined manually by calculating the ratio of nuclei in α-actinin-positive myotubes that contain more than two nuclei to the total number of nuclei.

### DNA constructs and virus production

Mouse *Slc25a34* cDNA was amplified from Mouse Untagged Clone (MC214787, ORIGENE) and ligated into a pcDNA3.1 plasmid (Thermo Fisher Scientific). For retrovirus production, HEK293T packaging cells were transfected with 10 µg of pMXs-3XHA-EGFP-OMP25, 5 µg of gag/pol, and 5 µg of VSV-G using the calcium phosphate method. After 48 hours of transfection, the virus-containing supernatant was collected, filtered through a 0.45 µm filter, and stored at -80°C for future use. Primary myoblasts were infected with the virus and polybrene (10 µg/mL) for 24 hours and then sorted with GFP using FACS.

### Mitochondrial isolation and metabolomics

Primary *Slc25a34*^flox/flox^ myoblasts stably expressing a mitochondria-localized epitope-tag (HA-MITO) (*53*) were differentiated into myotubes for 2 days and then infected with adenovirus Ad-eGFP (control) or Ad-iCre (*Slc25a34* KO) in a 15 cm dish. Mitochondrial metabolomics was performed 2 days after infection. Control and *Slc25a34* KO myotubes were washed twice with ice-cold PBS and collected with KPBS (136 mM KCl, 10 mM KH2PO4, pH 7.25) into the tubes and then centrifuged for 2 min at 1,000 x *g* at 4°C. The collected cells were resuspended in 1 mL KPBS, and 100 µL of the cell suspension was added to 500 µL of 100% MeOH for whole-cell metabolomic analysis. The cells were homogenized using a Potter-homogenizer with a PTFE pestle to obtain a mitochondrial fraction and then centrifuged for 2 min at 1,000 x g at 4°C. The supernatant containing the mitochondrial fraction was mixed with 200 µL of Pierce™ Anti-HA Magnetic Beads (Thermo Scientific) on an end-over-end rotator for 4 min at 4°C. The beads were collected using a magnet rack for 1 min and carefully washed three times with KPBS. The bead suspension was collected using a magnet, and the supernatant was removed. Then 80% MeOH (80 µL) was added to the beads, and the samples were placed at -80°C overnight for metabolite extraction. After incubation, the samples were mixed well by vortexing and centrifuged for 10 min at 21,000 x g at 4°C. The supernatant was transferred to a new tube, and this step was repeated once, and stored at -80°C until LC-MS analysis. Metabolomics data were acquired using a UHPLC system (Vanquish Horizon, Thermo Scientific) coupled to an Orbitrap mass spectrometer (Exploris 240, Thermo Scientific) as described previously (*6*).

### Structural prediction of SLC25A34

The structural models of SLC25A34 and SLC25A35 were obtained from the AlphaFold Protein Structure Database (https://alphafold.ebi.ac.uk/) (*37*) using the codes AF-Q6PIV7-F1-v6 for SLC25A34 and AF-Q3KQZ1-F1-v6 for SLC25A35. The predicted structures of SLC25A34 and SLC25A35 were aligned and visualized using UCSF ChimeraX-1.10.1 software (*80*). Protein sequence alignment and homology between human SLC25A34 and human SLC25A35 were analyzed using the EMBOSS Needle tool (*81*).

### Purification of MBP- and TwinStrep-tagged SLC25A34

Codon-optimized N-terminally MBP-tagged and C-terminally Twin-Strep-tagged human wild-type SLC25A34 or mutant SLC25A34 or SLC25A35 cDNA was synthesized as gBlock gene fragments (IDT) and inserted into the pET-21b(+) expression vector (V011022, NovoPro) using the In-fusion Cloning Kit (638948, Takara). Bacteria (C43(DE3)) transformed with pET-21b(+) empty or MBP-hSLC25A34-TwinStrep were cultured at 37°C for 6 hours in LB medium containing 100 µg/mL ampicillin, followed by a 1:100 dilution in 500 mL of LB medium containing 100 µg/mL ampicillin. After 1 hour of incubation at 37°C, IPTG (0.1 mM) was added to induce protein expression, followed by overnight induction (20°C, 300 rpm., 14 hours). After induction, bacteria were collected and washed once with PBS, and then resuspended in W buffer (2-1003-100, IBA) containing a protease inhibitor cocktail (Roche). The resuspended pellets were lysed by high-pressure homogenization using a French Press (27 kpsi). The lysate was centrifuged at 600 x g for 10 min to remove the debris and supplemented with 2% Triton X-100 and rotated at 4°C for 2 hours. The lysate was centrifuged at 21,000 x g for 15 min, and the supernatant was rotated with a Strep-TactinXT 4Flow high-capacity resin (2-5030-010, IBA) at 4°C for 2 hours. The resin was washed five times with W buffer containing 2% Triton X-100 and a protease inhibitor cocktail, followed by four rounds of rotation in BXT buffer (2-1042-025, IBA) containing 1% Triton X-100 and a protease inhibitor cocktail at 4°C for a total of 2 hours to obtain purified proteins.

### Bacterial PEP uptake by SLC25A34

Bacterial expression of SLC25A34 proteins and uptake assays were performed as described previously, with modification (*38, 82, 83*). MBP-tagged fusion human SLC25A34 was generated by combining *E. coli* MBP containing the MalE signal peptide and the *Slc25a34* gene. Each protein was fused via a thrombin-cleavage site. MBP and codon-optimized hSLC25A34 coding sequences were synthesized as gBlock gene fragments (IDT) and cloned into the pET-21b(+) expression vector. Expression of MBP-hSLC25A34 was achieved in *E. coli* C43(DE3) cells (CMC0019-20X40UL; Biosearch Technologies) (*84, 85*). According to the manufacturer’s protocol, single aliquots of competent bacterial cells were transformed with pET-21b(+) containing MBP-hSLC25A34 and selected on LB agar plates containing 100 µg/mL ampicillin (37°C, overnight). Bacteria transformed with pET-21b(+) empty or MBP-hSLC25A34 plasmids were cultured overnight in LB medium containing 100 µg/mL ampicillin, followed by a 1:100 dilution in 300 mL of LB medium containing 100 µg/mL ampicillin. Refreshed bacteria were incubated (37°C, 200 rpm) until OD600 reached 0.3-0.4, and the cultures were removed and cooled on ice. The prepared bacteria were split into 50 mL aliquots in 250 mL flasks. 0.1 mM isopropyl β-D-1-thiogalactopyranoside (IPTG, I56000, Research Products International) was added to the chilled bacteria to induce protein expression, followed by overnight induction (20°C, 300 rpm, 14 hours). MBP-SLC25A45-expressing bacteria were prepared as previously described (*40*). The protein expression of SLC25A34 or SLC25A45 in bacteria was confirmed by immunoblotting.

Induced bacteria were collected by centrifugation (4,000 x g, 10 min, 4°C), washed once with ice-cold potassium phosphate buffer (KPi), pH 7.4, and centrifuged again (4,000 x g, 15 min, 4°C). The bacteria were resuspended in ice-cold KPi buffer, and the OD600 was measured. Thirty OD600*mL units of bacteria transformed with empty vector or overexpressing MBP-hSLC25A34 were resuspended in 1 mL of KPi buffer. For the uptake assay, 100 µM ^13^C_2_-PEP (CLM-3398, Cambridge Isotope Laboratories) or 1 mM ^13^C_4_-OAA (O845032, Toronto Research Chemicals) was added to the sample and shaken at 37°C for 30 min. For time-course experiments, the samples were incubated with 100 µM ^13^C_2_-PEP, and aliquots were taken at 5, 15, 30, and 60 min. For competition assays, the samples were incubated with 100 µM ^13^C_2_-PEP and unlabeled (^12^C) 50 mM PEP at 37°C for 30 min. Samples were centrifuged at 20,000 x g for 1 min and washed twice with KPi buffer. Metabolites were extracted with ice-cold 80% MeOH (PEP) or 50% Acetonitrile/30% MeOH (OAA) for LC-MS analysis. Bacterial uptake of ^13^C-labeled metabolites was analyzed by LC-MS and normalized to the total protein content of each sample.

### Proteo-liposome assay

To prepare liposomes, 100 mg of the lipids (Egg PC, *E. coli* Polar lipids, 18:1 Cardiolipin at a 4:4.2:9 ratio, Avanti Polar Lipids) in 10 mL of chloroform was incubated in a rotary evaporator at 100 rpm at 50°C overnight. The lipid film on the internal surface of the flask was rehydrated with 2 mL of PIPES buffer (10 mM PIPES, 50 mM NaCl, pH 7.0) containing 20 mM unlabeled PEP. The liposomes were extruded 15 times using a mini-extruder with a 200-nm pore membrane at 60°C. Extruded liposomes were rotated with purified SLC25A34 protein or background proteins (purified from empty vector-transformed bacteria) at 4°C for 1 hour. Proteo-liposomes were then rotated with Bio-Beads SM-2 five times to completely remove Triton X-100. The resultant proteo-liposomes were isolated on a PD-10 desalting column (17-0851-01, Cytiva) to remove the external substrates and centrifuged at 100,000 rpm for 10 min. Proteo-liposomes were incubated with 5 mM ^13^C_2_-labeled PEP in PIPES buffer at 37°C for 20 min and centrifuged at 100,000 rpm for 5 min. The pellet was washed twice with PIPES buffer, followed by centrifugation at 100,000 rpm for 5 min. The final pellet was lysed in 80% MeOH for LC-MS analysis to detect ^13^C_2_-labeled PEP.

### Mitochondrial uptake assays

Control and *Slc25a34* KO myotubes were prepared by adenovirus infection in 10 cm dishes. The cells were collected in KPBS and centrifuged for 10 min at 600 x g at 4°C. The pellets were resuspended in KPBS and homogenized using a Potter-homogenizer with a PTFE pestle. Homogenates were centrifuged for 10 min at 600 x g at 4°C to remove cell debris and the mitochondria fraction containing supernatant was transferred into a new tube and then centrifuged for 10 min at 7,000 x g at 4°C. The pellet was incubated in KPBS containing 0.32 µCi/mL of 1,5-^14^C-Citrate, [U-^14^C]-Malic acid, 1-^14^C-α-ketoglutarate, [U-^14^C]-Aspartic acid or [U-^14^C]-Leucine (Moravek) for 1 hour at 4°C, or 1 mM of [U-^13^C]-α-ketoisocaproic acid (KIC), [U-^13^C]-α-ketoisovaleric acid (KIV), [U-^13^C]-α-keto-β-methylvaleric acid (KMV) (CIL) and 10 µM of ^13^C_2_-PEP for 5 min at 4°C. After incubation, ^14^C-containing mitochondria were washed twice with ice-cold KPBS and lysed with 500 µL of RIPA buffer. Radioactivity was measured using a scintillation counter and normalized to the total protein content. Metabolites were extracted using 80% MeOH for LC-MS analysis.

### Derivatization

To quantify ^13^C_4_-OAA, we used 3-nitrophenylhydrazine (3-NPH) to convert it into ^13^C_4_-malate following a published method (*43*). The extracted solvent containing ^13^C_4_-OAA was derivatized with 3-NPH (N21804, Sigma). The solvent was mixed with 200 mM 3-NPH in 50% acetonitrile and 120 mM N-(3-Dimethylaminopropyl)-Ń-ethylcarbodiimide (EDC, E1769, Sigma) in 50% acetonitrile/6% pyridine (1.09728, Sigma) and incubated at 4°C for 45 min with gentle shaking. After incubation, the samples were centrifuged for 5 min at 20,000 x g at 4°C and used for LC-MS analysis. All reagents were freshly prepared.

### Whole-cell tracing experiments

For glucose tracing, differentiated primary myotubes were incubated with 5.5 mM [U-^13^C]-Glucose (Cambridge Isotope Laboratories, CLM-1396-1) in HBSS (137 mM NaCl, 5.4 mM KCl, 0.34 mM Na2HPO4, 0.44 mM KH2PO4, 4.2 mM NaHCO3, 1.26 mM CaCl_2_ and 0.4 mM MgSO_4_) for 6 hours. For pyruvate and glutamine tracing, the cells were incubated for 6 hours in HBSS containing 5.5 mM glucose supplemented with either 2 mM [U-^13^C]-Pyruvate (Cambridge Isotope Laboratories, CLM-2440) or 2 mM [U-^13^C]-Glutamine (Cambridge Isotope Laboratories, CLM-1822). After incubation, the cells were washed twice with ice-cold PBS and collected for metabolite extraction with ice-cold 80% MeOH and stored at -80°C until LC-MS analysis. The labeling rate of each metabolite was calculated by dividing the level of labeled metabolites by the total metabolite level.

### Electron microscopy and quantification

: Soleus muscle was immersion fixed in 2.5% glutaraldehyde (Electron Microscopy Sciences, Hatfield, PA), 2% paraformaldehyde (Electron Microscopy Sciences), and 0.3% picric acid (Sigma-Aldrich, St. Louis, MO) in 0.1 M sodium cacodylate buffer (Sigma-Aldrich) pH 7.4 for 1 hour at room temperature then at 4°C overnight. Tissues were washed with 0.1 M sodium cacodylate buffer, and then post-fixed for 1 hour at 4°C in 1% osmium tetroxide (Electron Microscopy Sciences) in 0.1M sodium cacodylate buffer. Tissues were washed in DI water and incubated in 2% aqueous uranyl acetate (Electron Microscopy Sciences) overnight at 4°C. The following day, tissues were washed with DI water and then dehydrated at 4°C in a graded ethanol series (Thermo Fisher Scientific, Waltham, MA). The tissues were then brought to room temperature and dehydrated with 100% ethanol (Thermo Fisher Scientific) followed by propylene oxide (Electron Microscopy Sciences). Infiltration in LX112 resin (Ladd Research Industries, Williston, VT), was followed by embedding in flat bottom Beem capsules (Electron Microscopy Sciences). The resulting blocks were sectioned using a Leica Ultracut E ultramicrotome (Leica Biosystems Nussloch, Germany), and sections were placed on formvar (Electron Microscopy Sciences) and carbon-coated grids. The sections were contrast stained with 2% uranyl acetate (Electron Microscopy Sciences) followed by lead citrate (Sigma-Aldrich) and imaged in a JEOL 1400 transmission electron microscope (JEOL, Peabody, MA) equipped with a Gatan Orius SC1000 digital CCD camera (Gatan, Pleasanton, CA). The mitochondrial area was quantified by tracing the outer membrane using a free-hand tool and normalizing to scale.

### Respiration in isolated muscle fiber

A portion of the soleus and plantaris muscles was dissected and placed in Buffer Z (1 mM EGTA, 5 mM MgCl_2_, 105 mM K-MES, 30 mM KCl, 10 mM KH2PO4, 5 mg/mL fatty acid-free BSA, pH 7.2). Muscle fibers were separated under a microscope and permeabilized in Buffer Z with saponin (30 µg/mL) at 4°C for 30 min. The muscle fibers were incubated in ice-cold Buffer Z with pyruvate (5 mM)/malate (2 mM) and then transferred to the chamber of an Oroboros O2k respirometer (Oroboros Instruments) in Buffer Z. Respiration was measured in Buffer Z with 5 mM pyruvate, 2 mM malate, 0.2 mM ADP, 5 mM succinate, and 5 µM rotenone. After the experiments, the muscle fibers were washed, and the protein concentration was measured to normalize the respiration data.

### Cellular respiration assay

Oxygen consumption rate (OCR) and extracellular acidification rate (ECAR) were measured using the Seahorse XFe Extracellular Flux Analyzer (Agilent) in a 24-well plate. Control and *Slc25a34* KO primary myotubes were cultured in standard assay medium containing 25 mM glucose, 2 mM glutamine and 1 mM pyruvate. For the measurement of uncoupled respiration, myotubes were treated with 2 µM oligomycin, 1.5 µM carbonyl cyanide-p-trifluoromethoxyphenylhydrazone (FCCP) and 2 µM antimycin A.

### Immunofluorescent imaging

HEK293T cells were placed in a glass-bottom dish (10810-054, VWR) and cultured for 24 hours. The cells were then transfected with 100 ng of pcDNA3.1-SLC25A34 plasmid using Lipofectamine 3000 (Thermo Fisher Scientific). 24 hours after transfection, the cells were washed twice with PBS and fixed with 4% paraformaldehyde (PFA) at 37°C for 30 min. The cells were washed three times with PBS and permeabilized with 0.3% NP-40, 0.05% Triton X-100, and 0.1% bovine serum albumin (BSA) in PBS for 3 min. After three rinses with wash buffer (0.05% NP-40, 0.05% Triton X-100, and 0.2% BSA in PBS), the samples were blocked for 1 hour at room temperature with SuperBlock Blocking Buffer (37515, Thermo Fisher Scientific). The samples were then incubated with primary antibodies in wash buffer overnight at 4°C, washed three times with wash buffer. The primary antibodies used targeted TOMM20 (11802-AP, Proteintech) and FLAG (8146S, Cell Signaling Technology). Secondary antibodies were conjugated with Alexa Fluor 488 (ab150117, abcam) and Alexa Fluor 647 (A21245, Invitrogen). Images were acquired using Zeiss LSM900 confocal microscope with Airyscan and processed using Zeiss Zen software.

Control and *Slc25a34* KO myotubes were fixed with 4% PFA at 37°C for 10 min, washed with PBS, and then permeabilized with 0.1% Triton X-100 in PBS at room temperature for 15 min. The cells were then washed with PBS and blocked with 2% BSA in PBS at room temperature for 1 hour and incubated with the α-actinin antibody (A7811, Sigma) at room temperature for 1 hour. After washing with PBS, the cells were incubated with Alexa Fluor 594-conjugated anti-mouse IgG1 secondary antibody and DAPI at room temperature for 1 hour. Images of the myotubes were acquired using the Revolve microscope (ECHO Laboratories).

Frozen cross-sections from OCT-embedded muscle tissues were fixed with 4% PFA in PBS and washed once with PBS. The samples were permeabilized with 0.3% Triton X-100 in PBS, washed twice with PBS and then blocked with 5% normal goat serum in PBS. The tissue sections were then incubated at 4°C overnight with the following primary antibodies: Laminin (L9393, Sigma), myosin heavy chain (MyHC) I (BA-F8, DSHB), MyHC IIa (SC-71, DSHB), or dystrophin (D8043, Sigma). The secondary antibodies were Alexa Fluor 647-conjugated anti-rabbit IgG for laminin, DyLight 405-conjugated anti-mouse IgG2b for MyHC I, Alexa Fluor 488-conjugated anti-mouse IgG1 for MyHC IIa and dystrophin (Jackson ImmunoResearch Lab). Images were acquired using the Revolve microscope (ECHO Laboratories).

### Immunoblotting

Skeletal muscle tissues were homogenized in complete protein loading buffer containing 50 mM Tris-HCl (pH 6.8), 1% SDS, 10% Glycerol, 20 mM dithiothreitol (DTT), 127 mM 2-mercaptoethanol, and 0.01% bromophenol blue, supplemented with protease inhibitors and phosphatase inhibitors (Sigma). The protein contents were measured with an RC DC Protein Assay Kit (Bio-Rad) according to the manufacturer’s instructions. The proteins were electrophoresed on SDS-PAGE precast gels (Bio-Rad) and transferred to a PVDF membrane (Millipore), and the signals were immunodetected with Clarity Western ECL Substrate (Bio-Rad) using the ChemiDoc Imaging System (Bio-Rad). The following antibodies were used: Puromycin (MABE343, Sigma), S6K (#2708, CST), p-S6K (#9205, CST), p-rp-S6 (#5364, CST), rp-S6 (#2217, CST), Strep-Tactin^®^-HRP (2-1502-001, IBA), MBP tag (66003-1-Ig, Proteintech), ATP5A (ab14748, Abcam), Total OXPHOS (ab110413, Abcam), AMPK alpha (#2532, CST), p-AMPK alpha (#2535, CST) and GAPDH (2118S, CST). Anti-Rabbit IgG, HRP-Linked antibody (ab6721, Abcam) and anti-mouse IgG HRP (#31430, Thermo Fisher Scientific) were used as secondary antibodies.

### Animal physiology

The endurance exercise capacity test was performed as described previously (*86*). Mice were acclimatized to the treadmill for 5 min at 7 m/min on a 10% incline on two consecutive days. The test began at 8 m/min for 30 min, followed by 9 m/min for 15 min and 10 m/min for 15 min. The speed was then increased by 1 m/min every 10 min until the mice were exhausted. Exhaustion was defined as the point at which the mice were unable to return to the treadmill belt after 10 sec of encouragement. Grip strength was measured using a grip strength meter (Ametek) by placing all four limbs of the mice on a metal grid. The peak force was recorded in each of five trials, and the median value was used for analysis.

### SUnSET assay

*In vivo* SUnSET assay was performed to evaluate nascent protein synthesis as described previously (*63*). Mice were intraperitoneally injected with puromycin (0.04 µmol/g body weight) dissolved in 100 µL PBS. At precisely 30 min after injection, soleus and plantaris muscles were dissected and immediately frozen in liquid nitrogen for immunoblotting. To inhibit mTORC1 signaling, mice were intraperitoneally injected with rapamycin (1.5 mg/kg) for 7 consecutive days.

### RNA-sequencing and Data Analysis

Total RNA was purified from the soleus using the RNeasy® Fibrous Tissue Mini Kit (Qiagen) with DNase treatment on the column. Total RNA samples were quantified using a Qubit 2.0 Fluorometer (Life Technologies), and RNA integrity was checked using a 4200 TapeStation (Agilent Technologies). The samples were initially treated with TURBO DNase (Thermo Fisher Scientific) to remove DNA contaminants. ERCC RNA Spike-In Mix (Thermo Fisher) was added to normalized total RNA prior to library preparation, following the manufacturer’s protocol. The next steps included rRNA depletion using the QIAGEN FastSelect rRNA HMR Kit (Qiagen). RNA sequencing libraries were multiplexed and clustered on the flowcell. After clustering, the flowcell was loaded onto an Illumina NovaSeq Instrument. The samples were sequenced using a 2x150 Paired-End (PE) configuration. After demultiplexing, the sequence data were checked for overall quality and yield. Then, raw sequence reads were trimmed to remove possible adapter sequences and nucleotides with poor quality using Trimmomatic v.0.36. The reads were then mapped to the reference genome (mm10) using the STAR aligner v.2.5.2b. Unique gene hit counts were calculated by using featureCounts from the subread package v.1.5.2. Only unique reads that fell within exon regions were counted. Downstream analysis was performed in R. Differential expression analysis was performed using DESeq2. P values were calculated using the Wald test and adjusted for multiple testing using the Benjamini-Hochberg procedure. Genes with Benjamini-Hochberg-adjusted p value (false discovery rate; q value) < 0.05 were considered differentially expressed. Gene enrichment analysis was performed using Enrichr (*87*).

### RT-qPCR

Total RNA was isolated from cells and non-muscle tissues using the Direct-zol RNA Purification Kit (Zymo Research), and from skeletal muscle using the RNeasy® Fibrous Tissue Mini Kit (Qiagen), according to the manufacturer’s protocols. For the measurement of mRNA expression, 1 µg of total RNA was reverse transcribed with Oligo-dT(20) primer using SuperScript™ IV Reverse Transcriptase (Thermo Fisher Scientific). The synthesized cDNA and SsoAdvanced Universal SYBR® Green Supermix (Bio-Rad) were used to quantify mRNA levels. GAPDH and 18S rRNA were used as the internal controls. The primer sequences are presented in **Supplementary Table 10**.

### Statistics

Statistical analyses were conducted using GraphPad Prism version 10 (GraphPad). All data are represented as mean ± s.e.m. Data were obtained from biologically independent samples. *P* < 0.05 was considered to be significant throughout the study. For comparison between two groups, unpaired Student’s *t*-tests or the Mann-Whitney test were employed. To compare three or more groups, one-way ANOVA followed by suitable post-hoc tests was applied. To assess the effects of two independent factors, two-way ANOVA or two-way repeated-measures ANOVA was applied with Tukey’s multiple comparisons test. For metabolomics and SLC25A family transcriptomics analyses, unpaired *t*-tests followed by Benjamini-Hochberg FDR correction were used.

## Acknowledgments

We are grateful to Naofumi Yoshida, Christopher Auger, Kazusa Miyachi, Yusuke Higuchi, Alice Ma, and Mia Formato in the Kajimura Lab for their technical support. We also thank Dr. Suzanne L White at Beth Israel Deaconess Medical Center for her support in tissue histology study, and the EM core facility at Beth Israel Deaconess Medical Center for support in electron microscopy study.

## Author contributions

Conceptualization: S.O., S.K. Methodology: S.O., T.Y., H.N., D.K., D.W. Investigation: S.O., T.Y., H.N., D.K., D.W., M.F. Formal Analysis: S.O., H.N., D.W., M.F. Visualization: S.O., H.N., M.F. Funding acquisition: S.K. Supervision: S.K. Writing – original draft: S.O., S.K. Writing – reviewing & editing: all authors.

## Data, code, and materials availability

The raw data of metabolomics by LC-MS are uploaded to the Metabolomics Workbench (https://www.metabolomicsworkbench.org), with a project ID PR002784 and a project DOI: http://dx.doi.org/10.21228/M8SV79. The raw and processed data of RNA-seq generated in this study have been deposited in the Gene Expression Omnibus (GEO) database under accession code GSE311513. Previously published datasets used in this study include: GSE210904, GSE123879, GSE120862, GSE51178, GSE221210, GSE162455, GSE3307, GSE226646, GSE100505, GSE70437 and GSE134962. All other data needed to evaluate and reproduce the results in the paper are present in the paper and/or the Supplementary Materials. All the materials and mice are available after completion of a material transfer agreement.

---

**Figure S1.**
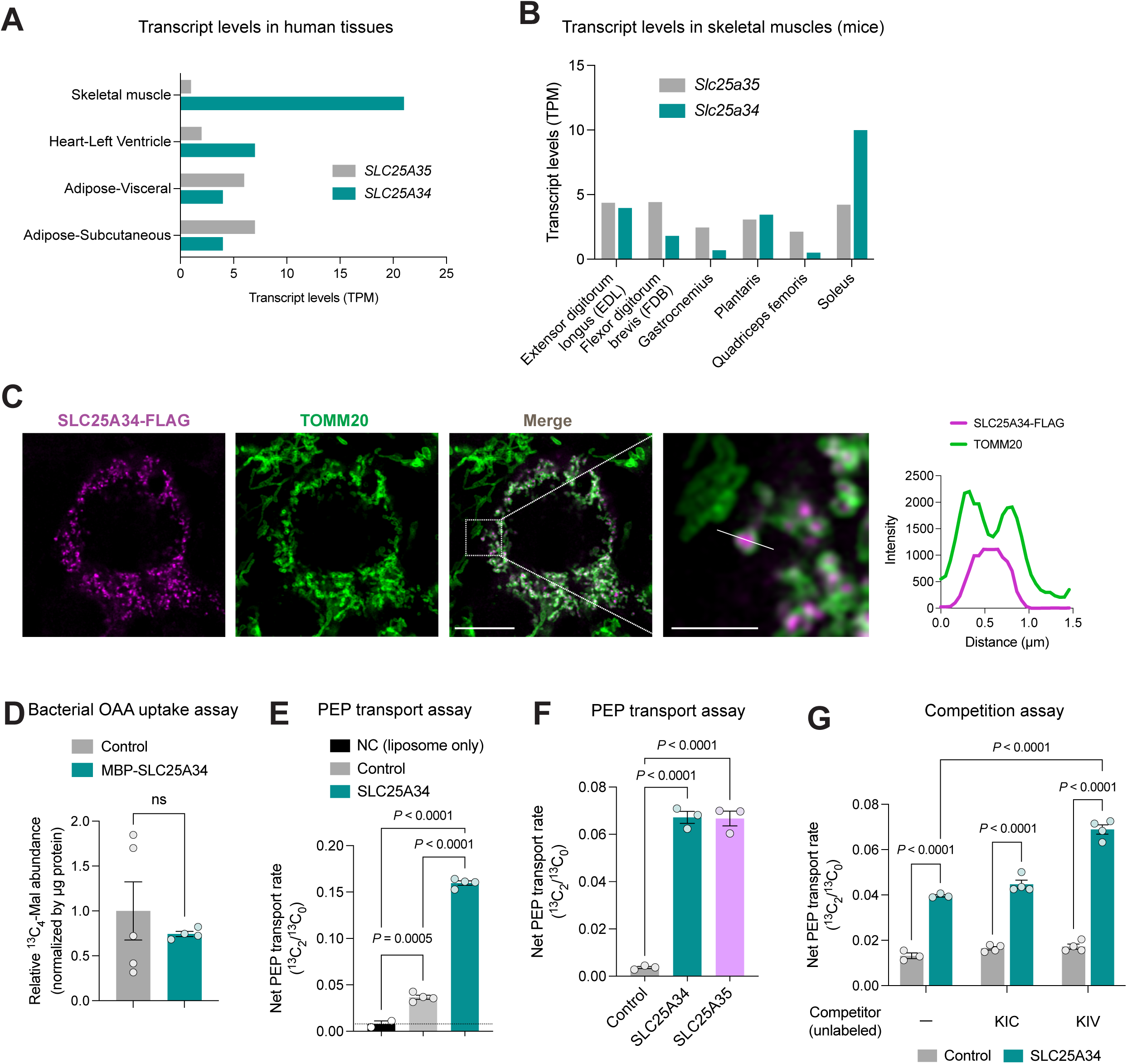
Molecular characterization of SLC25A34, related to Figure 1. **A.** Transcript levels (TPM) of *SLC25A34* and *SLC25A35* in the indicated human tissues. Data were obtained from GTEx Portal (https://www.gtexportal.org/home/). **B.** Transcript levels (TPM) of *Slc25a34* and *Slc25a35* in the extensor digitorum longus (EDL), flexor digitorum brevis (FDB), gastrocnemius, plantaris, quadriceps femoris, soleus (GEO: GSE100505). **C.** Representative immunofluorescence image of HEK 293T cells with SLC25A34-Flag (magenta) and TOMM20 (mitochondrial outer membrane; green). Scale bars, 20 μm. The inset shows higher-magnification views of the boxed regions. Scale bar, 5 µm. The graph panel shows the fluorescence intensity along the solid white line. **D.** Bacterial oxaloacetate (OAA) uptake assay. Bacteria expressing the endogenous MBP protein alone (control) or MBP-SLC25A34 were incubated with ^13^C_4_-OAA (1 mM) for 30 min at 37°C. Derivatized ^13^C_4_-malate, a downstream metabolite of OAA, was analyzed by LC-MS and normalized to the protein levels in each sample. n = 5 for control and n = 4 for MBP-SLC25A34. **E.** PEP transport assay in proteo-liposomes reconstituted with purified SLC25A34 or control-eluates. Background signals obtained from protein-free empty control liposomes (*i.e.,* liposome lipids only) were subtracted, as these signals represent non-specific association with the liposomes. n = 2 for NC (liposome lipids only), n = 4 for control and SLC25A34. **F.** PEP transport assay in proteo-liposomes reconstituted with purified SLC25A34 or SLC25A35 or control-eluates. n = 3 per group. **G.** Competition assay. An excess amount of indicated metabolites (50 mM) was added to the assay. n = 3 for non-competitor and n = 4 for KIC and KIV. Bars represent mean ± s.e.m. *P* values were calculated by unpaired *t*-test (D), one-way ANOVA with Tukey’s multiple comparisons test (E and F) and two-way ANOVA with Tukey’s multiple comparisons test (G).

**Figure S2.**
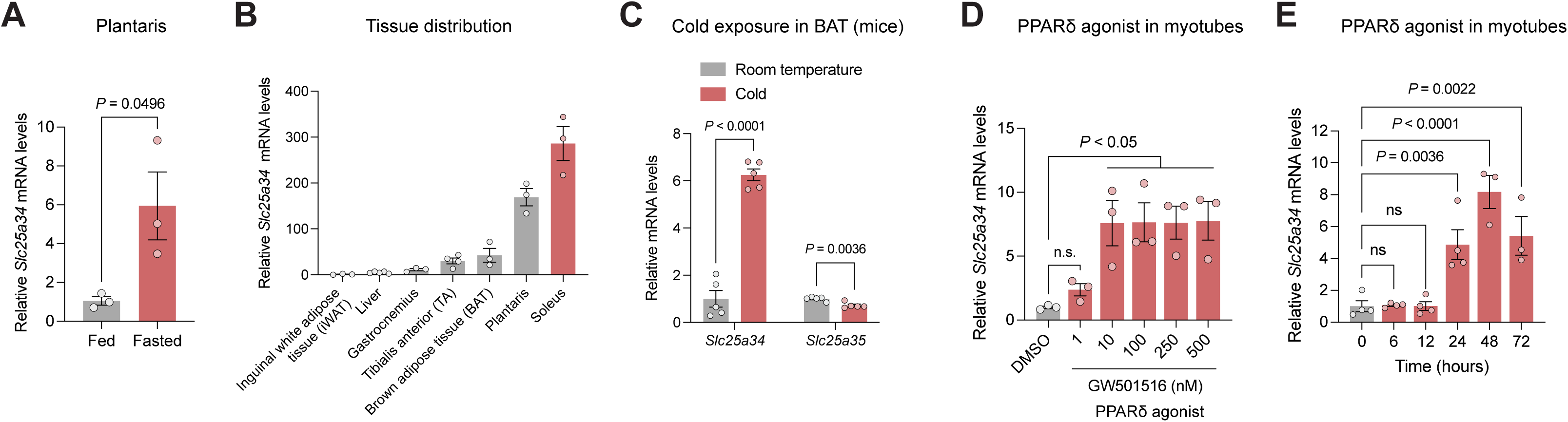
Regulation of SLC25A34 expression, related to Figure 2. **A.** Relative mRNA levels of *Slc25a34* in plantaris muscles from wild-type mice fasted for 24 hours, measured by qPCR. n = 3 per group. **B.** Relative mRNA levels of *Slc25a34* in the indicated tissues from wild-type mice, as measured by qPCR. n = 3 for inguinal white adipose tissue (iWAT), gastrocnemius, brown adipose tissue (BAT), plantaris, soleus; n = 4 for tibialis anterior (TA); n = 5 for liver. **C.** Relative mRNA levels of *Slc25a34* and *Slc25a35* in brown adipose tissue (BAT) from mice that were kept at room temperature (RT) or exposed to cold (4°C) for 3 days (GEO: GSE70437). n = 5 per group. **D.** Dose-dependent changes in *Slc25a34* expression levels upon PPARδ agonist treatment. Differentiated C2C12 myotubes were treated with the indicated concentrations of GW501516 for 24 hours. Relative *Slc25a34* mRNA levels were measured using qPCR. n = 3 per group. **E.** Time-dependent changes in *Slc25a34* expression levels following PPARδ agonist treatment. Differentiated C2C12 myotubes were treated with GW501516 (100 nM) for the indicated times (0-72 hours). Relative *Slc25a34* mRNA levels were measured using qPCR. n = 4 for 0, 6, 12, 24 hours and n = 3 for 48 and 72 hours. Bars represent mean ± s.e.m. *P* values were calculated by unpaired *t*-test (A and C) and one-way ANOVA with Dunnett’s multiple comparisons test (D and E).

**Figure S3.**
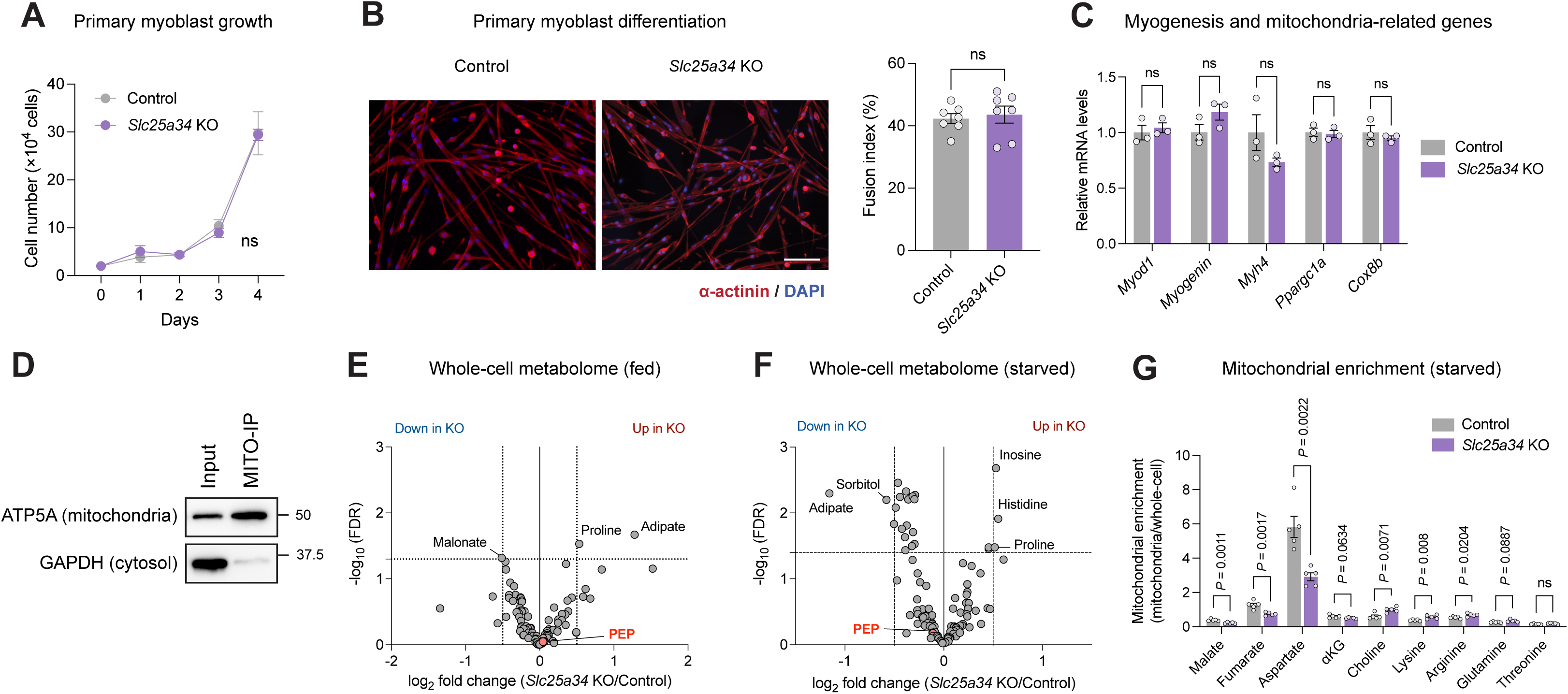
Myogenesis and metabolomics in *Slc25a34* KO primary myotubes, related to Figure 3. **A.** Growth of control and *Slc25a34* KO primary myoblasts. n = 4 at all time points. **B.** Representative immunofluorescence images of control and *Slc25a34* KO primary myotubes stained for α-actinin and DAPI. The fusion index was manually determined by calculating the ratio of nuclei in α-actinin-positive myotubes containing more than two nuclei to the total number of nuclei. n = 7 per group. **C.** Relative expression levels of myogenesis- and mitochondria-related marker genes in control and *Slc25a34* KO primary myotubes. n = 3 per group. **D.** Mitochondria purification using the MITO-Tag system. Mitochondria were isolated from primary myotubes expressing MITO-Tag (3xHA-EGFP-OMP25) using magnetic beads and subjected to immunoblotting to detect ATP5A (a mitochondrial marker) and GAPDH (a cytosolic marker). **E.** Whole-cell metabolomics of control or *Slc25a34* KO primary myotubes under a nutrient-replete culture condition (25 mM glucose, 5% horse serum, and amino acids). n = 5 per group. **F.** Whole-cell metabolomics of control or *Slc25a34* KO primary myotubes under a nutrient-deprived culture condition (HBSS containing 5.5 mM glucose). n = 5 per group. **G.** Mitochondrial metabolite enrichment in control or *Slc25a34* KO primary myotubes cultured in a nutrient-deprived condition. Mitochondrial enrichment was calculated as the ratio of mitochondrial to whole-cell intensity of the indicated metabolites. n = 5 per group. Bars represent mean ± s.e.m. *P* values were calculated by two-way ANOVA with Tukey’s multiple comparisons test (A), unpaired *t*-test (B, C, G) and unpaired *t*-test with Benjamini-Hochberg FDR correction (E and F).

**Figure S4.**
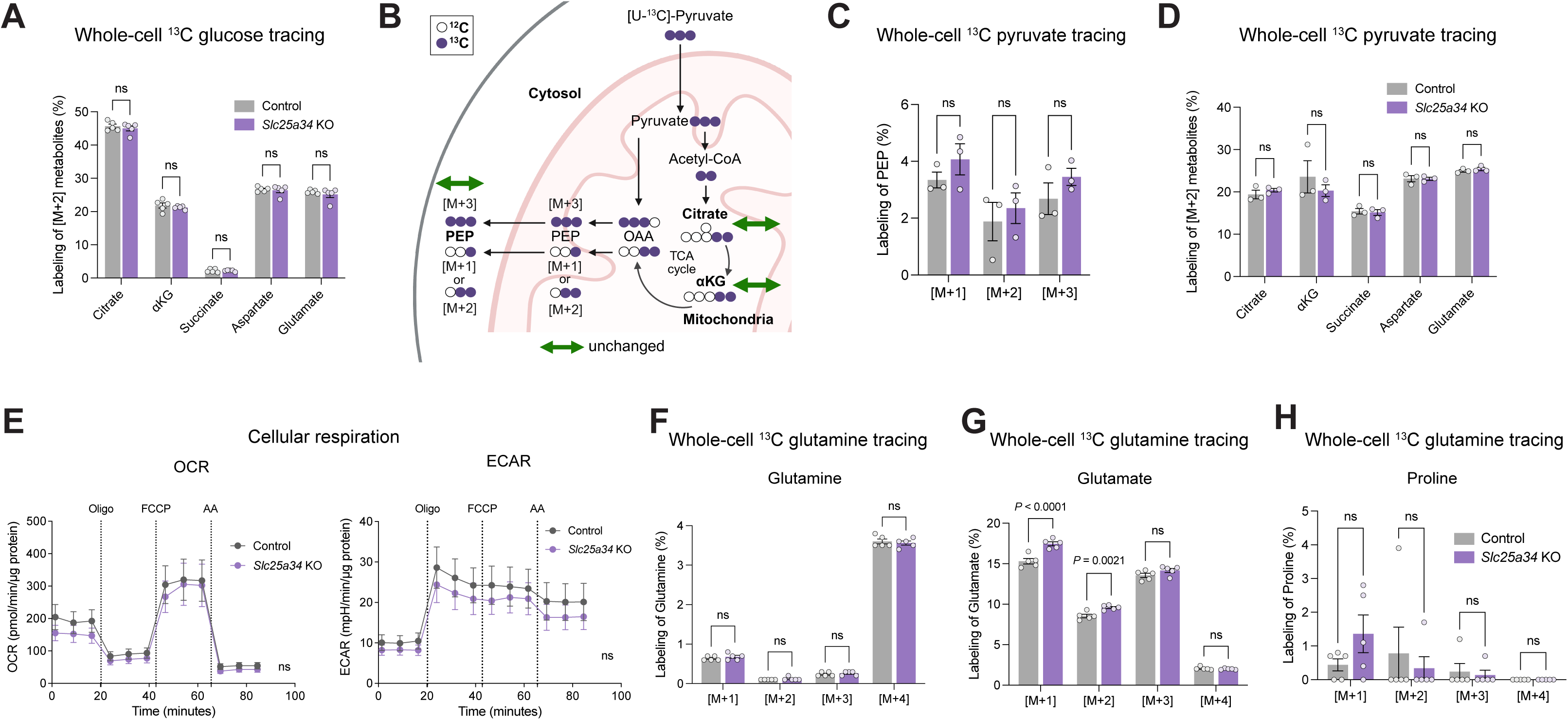
Stable isotope studies in *Slc25a34* KO primary myotubes, related to Figure 3. **A.** Whole-cell [U-^13^C]-glucose tracing in control or *Slc25a34* KO primary myotubes under a nutrient-deprived cultured condition (HBSS). The labeling of the indicated ^13^C-labeled [M+2] metabolites (%) is shown on the Y-axis. n = 5 per group. **B.** Schematic of [U-^13^C]-pyruvate whole-cell tracing. Control or *Slc25a34* KO primary myotubes were cultured in a nutrient-deprived medium with HBSS containing [U-^13^C]-pyruvate (1 mM) (green arrow, no change). **C.** Whole-cell [U-^13^C]-pyruvate tracing in control or *Slc25a34* KO primary myotubes cultured in a nutrient-deprived medium. The labeling of the indicated ^13^C-labeled PEP (%) is shown on the Y-axis. N = 3 per group. **D.** Whole-cell [U-^13^C]-pyruvate tracing in control or *Slc25a34* KO primary myotubes in a nutrient-deprived medium, HBSS containing 1 mM [U-^13^C]-pyruvate. The labeling of the indicated ^13^C-labeled [M+2] metabolites (%) is shown on the Y-axis. n = 3 per group. **E.** Cellular respiration in control and *Slc25a34* KO primary myotubes. Cellular oxygen consumption rate (OCR) and extracellular acidification rate (ECAR) were measured using the Seahorse XFe Extracellular Flux Analyzer. n = 9 per group. **F.** [U-^13^C]-glutamine tracing in control or *Slc25a34* KO cells under a nutrient-deprived medium condition. The labeling of the indicated ^13^C-labeled glutamine (%) is shown. n = 5 per group. **G.** [U-^13^C]-glutamine tracing in (F). The labeling of the indicated ^13^C-labeled glutamate (%) is shown. n = 5 per group. **H.** [U-^13^C]-glutamine tracing in (F). The labeling of the indicated ^13^C-labeled proline (%) is shown. n = 5 per group. Bars represent mean ± s.e.m. *P* values were calculated by unpaired *t*-test (A and D), two-way ANOVA with Tukey’s multiple comparisons test (C, F, G and H) and two-way repeated-measures ANOVA with Tukey’s multiple comparisons test (E).

**Figure S5.**
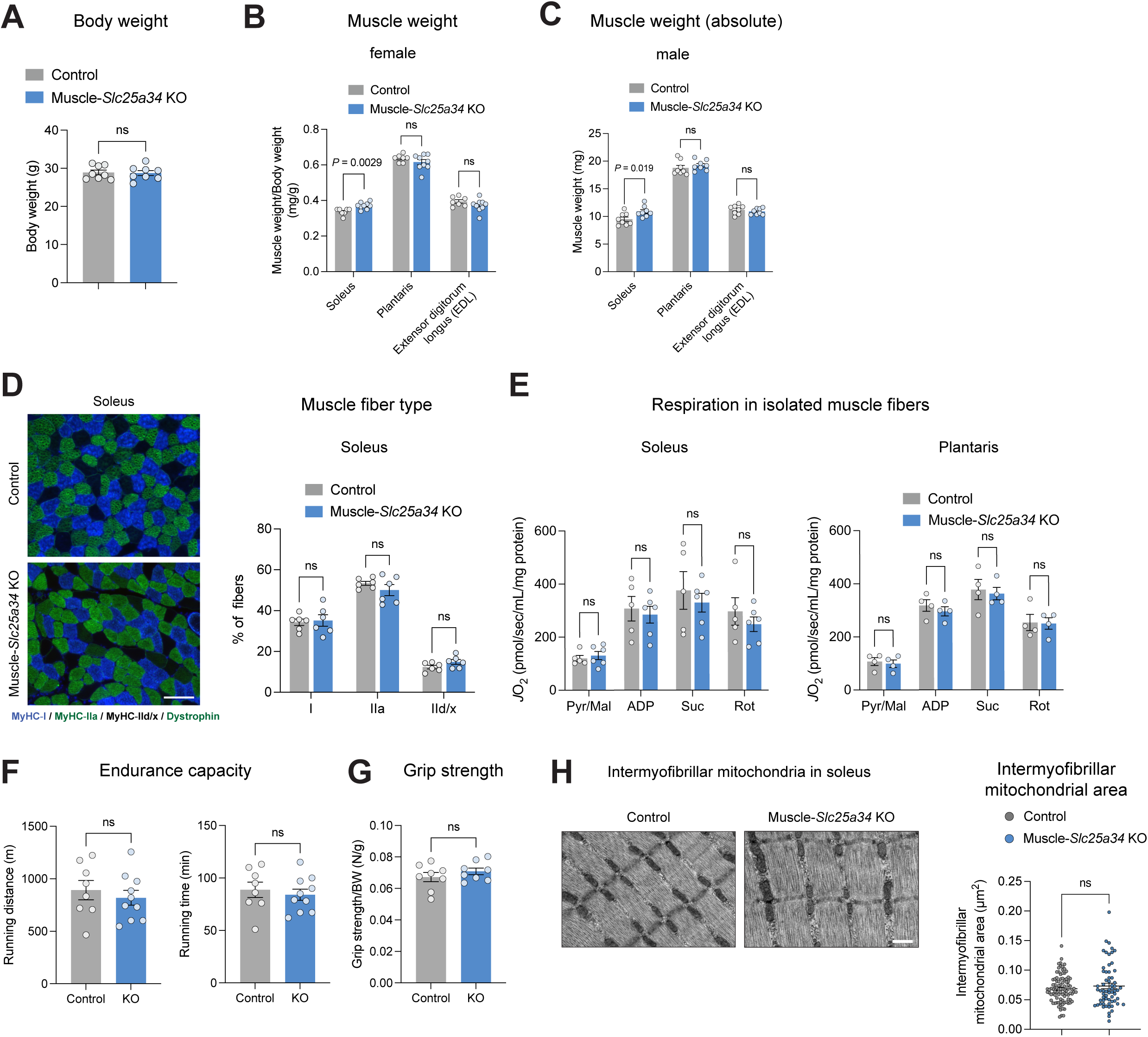
Characterization of muscle-*Slc25a34* KO mice, related to Figure 4. **A.** Body weight in male control and muscle-*Slc25a34* KO mice fed a regular diet at 16 weeks old. n = 8 per group. **B.** Muscle weight of the soleus, plantaris, and EDL in female control and muscle-*Slc25a34* KO mice at 17 weeks old. Muscle weight (mg) was normalized to the body weight (g). n = 7 for control and n = 9 for KO mice. **C.** Absolute muscle weight of soleus, plantaris, and EDL in male control and muscle-*Slc25a34* KO mice at 16 weeks old. n = 8 per group. **D.** Muscle fiber typing in soleus from male control and muscle-*Slc25a34* KO mice. n = 6 per group. **E.** Respiratory function in isolated muscle fibers from soleus (left) and plantaris (right) of male control and KO mice. n = 5 for control and n = 6 for KO in soleus, n = 4 per group for plantaris. **F.** Endurance performance in control and muscle-*Slc25a34* KO mice. Endurance exercise capacity assessed by running distance (left) and time (right) in male control and muscle-*Slc25a34* KO mice. n = 8 for control and n = 10 for KO. **G.** Four-limb grip strength in male control and muscle-*Slc25a34* KO mice. n = 8 per group. Grip strength (N) was normalized to the body weight (g). **H.** Representative electron microscopy images of intermyofibrillar mitochondrial structures in the soleus of male control and muscle-*Slc25a34* KO mice at 18 weeks old. Right: Quantification of intermyofibrillar mitochondrial area. Scale bar, 2 µm. n = 92 for control and 63 for KO collected from n = 2 mice per group. Bars represent mean ± s.e.m. *P* values were calculated by unpaired *t*-test (A, B, C, D, F, G and H) and two-way repeated-measures ANOVA with Tukey’s multiple comparisons test (E).

**Figure S6.**
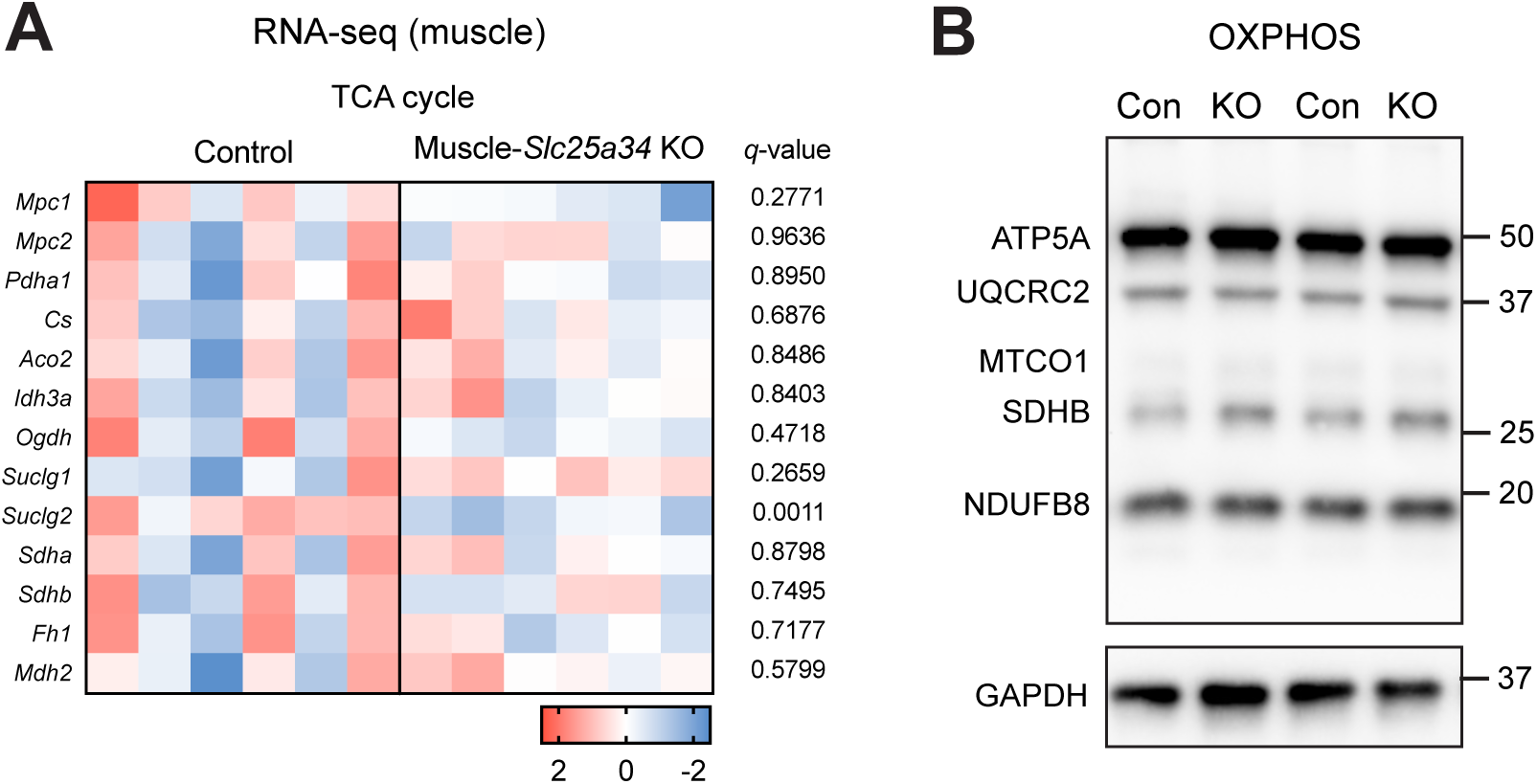
Molecular characterization of muscle-*Slc25a34* KO mice, related to Figure 5. **A.** Relative mRNA levels of TCA cycle-related genes in the soleus muscles of male control and muscle-*Slc25a34* KO mice at 18 weeks old. Data represented as *Z*-score heatmaps for each gene in each sample, based on RNA-seq analysis. n = 6 per group. Red indicates elevated expression and blue indicates decreased expression. **B.** Oxidative phosphorylation (OXPHOS) protein levels in the soleus muscle of male control and muscle-*Slc25a34* KO mice. Immunoblotting was performed to detect OXPHOS components (ATP5A, UQCR2, MTCO1, SDHB and NDUFB8) and GAPDH. *q*-values were calculated by Wald test with Benjamini-Hochberg FDR correction (A).

**Figure S7.**
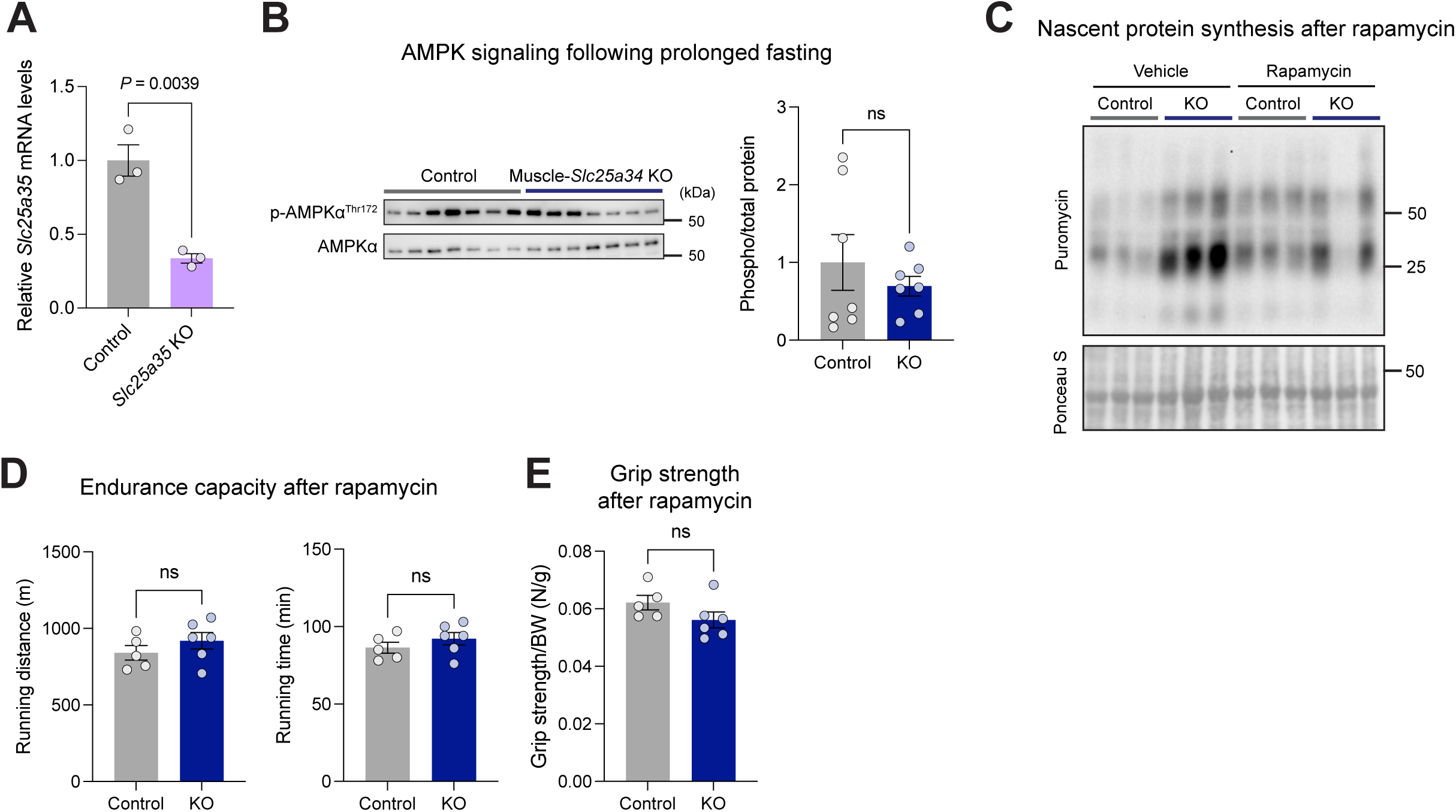
Molecular and physiological analyses of muscle-*Slc25a34* KO mice, related to Figure 6. **A.** Relative mRNA levels of *Slc25a35* in control and *Slc25a35* KO primary myotubes. *Slc25a35*^flox/flox^ cells were differentiated for 2 days, infected with adenovirus-GFP (Control) or adenovirus-Cre (KO), and analyzed by qPCR 2 days later. n = 3 per group. **B.** AMPK signaling in the soleus muscle after 24 hours of fasting. Immunoblotting was performed to detect phosphorylated-AMPK (Thr172). Phosphorylated protein levels were normalized to the corresponding total protein levels. n = 7 per group. **C.** Representative immunoblot of puromycin-labeled proteins in the soleus muscles of male control and muscle-*Slc25a34* KO mice following rapamycin treatment, assessed by the SUnSET assay. The total protein content (Ponceau S staining) was used as a loading control. **D.** Endurance performance in control and muscle-*Slc25a34* KO mice following rapamycin treatment. Endurance exercise capacity assessed by running distance (left) and running time (right) in male control and muscle-*Slc25a34* KO mice. n = 5 for control and n = 6 for muscle-*Slc25a34* KO. **E.** Four-limb grip strength in male control and muscle-*Slc25a34* KO mice following rapamycin treatment. n = 5 for control and n = 6 for muscle-*Slc25a34* KO. Grip strength (N) was normalized to the body weight (g). Bars represent mean ± s.e.m. *P* values were calculated by unpaired *t*-test (A, B, D and E).

**Supplementary Table 1.** Differential mitochondrial metabolites in control and *Slc25a34* KO primary myotubes under a fed condition (*P-*value < 0.05), related to Figure 3.

| Metabolite | FC | Log<sub>2</sub>(FC) | *P*-value | -log<sub>10</sub>(FDR) |
| --- | --- | --- | --- | --- |
| Guanosine | 2.1291 | 1.090217 | 0.004156 | 0.351972 |
| Adipate | 0.6224 | -0.68407 | 0.004525 | 0.616058 |
| N-Acetylaspartate | 1.4307 | 0.51675 | 0.004686 | 0.776947 |
| Citrate | 0.6995 | -0.51565 | 0.005481 | 0.833824 |
| Choline | 1.6641 | 0.73476 | 0.013059 | 0.553672 |
| Cytidine | 1.5089 | 0.593514 | 0.016549 | 0.529991 |
| Uridine | 2.0105 | 1.007585 | 0.019204 | 0.532312 |
| Oxalate | 0.7853 | -0.34864 | 0.022153 | 0.52828 |
| Inosine | 1.6680 | 0.738149 | 0.02789 | 0.479405 |
| Glutathione reduced | 2.1308 | 1.091385 | 0.028267 | 0.519336 |
| Malonate | 0.8566 | -0.22334 | 0.034897 | 0.469225 |
| Carnosine | 1.5247 | 0.608488 | 0.036524 | 0.48722 |
| Fad | 1.3527 | 0.43581 | 0.037001 | 0.516344 |
| N-Acetylmethionine | 1.4299 | 0.515932 | 0.038264 | 0.53395 |
| N-Acetylglutamate | 1.3246 | 0.40558 | 0.041589 | 0.527732 |
| S-Adenosylhomocysteine | 2.2248 | 1.153702 | 0.04203 | 0.551177 |
| Acetoacetate | 0.8150 | -0.29508 | 0.046787 | 0.530941 |

**Supplementary Table 2.**
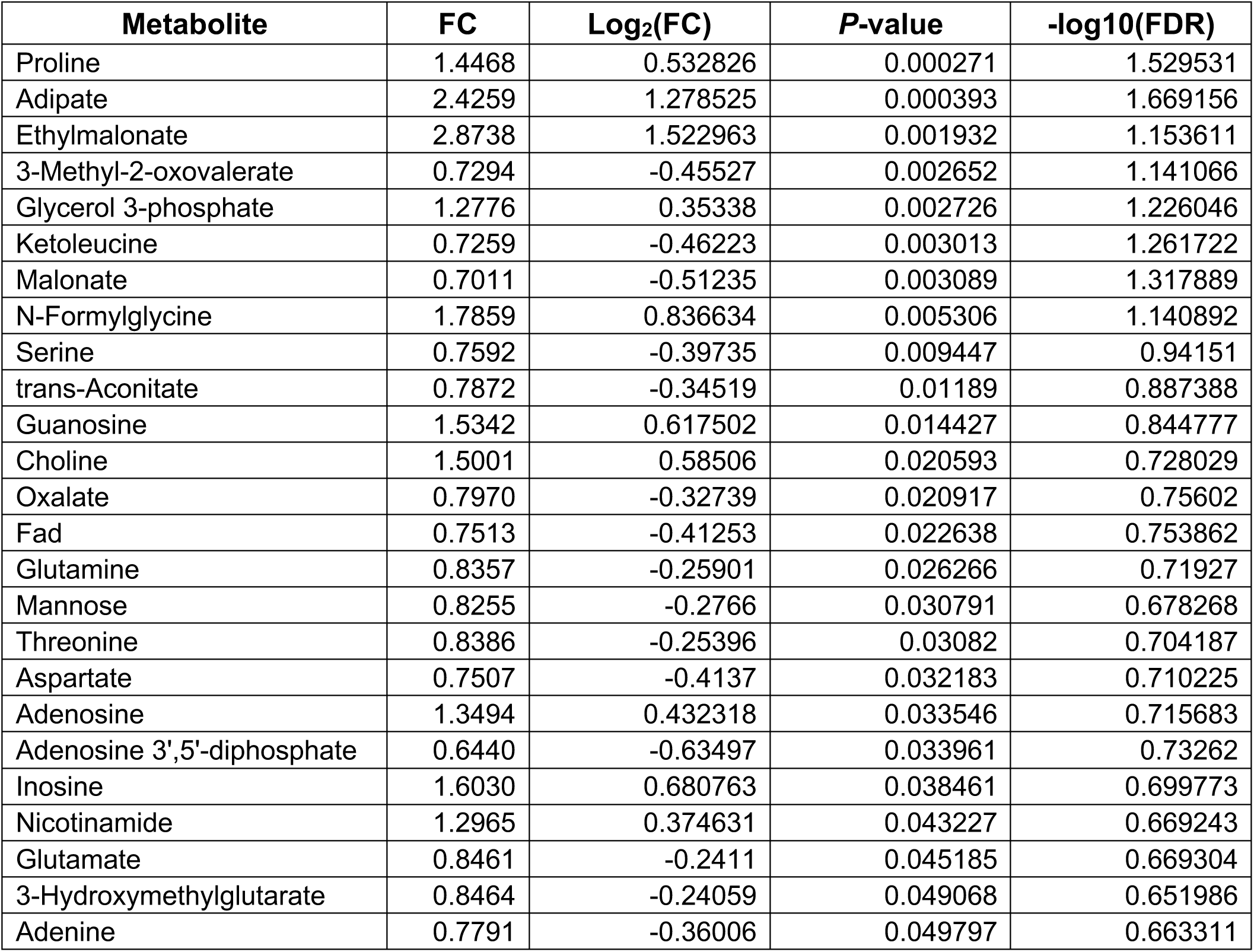
Differential whole-cell metabolites in control and *Slc25a34* KO primary myotubes under a fed condition (*P-*value < 0.05), related to Figure S3.

| Metabolite | FC | Log<sub>2</sub>(FC) | *P*-value | -log<sub>10</sub>(FDR) |
| --- | --- | --- | --- | --- |
| Proline | 1.4468 | 0.532826 | 0.000271 | 1.529531 |
| Adipate | 2.4259 | 1.278525 | 0.000393 | 1.669156 |
| Ethylmalonate | 2.8738 | 1.522963 | 0.001932 | 1.153611 |
| 3-Methyl-2-oxovalerate | 0.7294 | -0.45527 | 0.002652 | 1.141066 |
| Glycerol 3-phosphate | 1.2776 | 0.35338 | 0.002726 | 1.226046 |
| Ketoleucine | 0.7259 | -0.46223 | 0.003013 | 1.261722 |
| Malonate | 0.7011 | -0.51235 | 0.003089 | 1.317889 |
| N-Formylglycine | 1.7859 | 0.836634 | 0.005306 | 1.140892 |
| Serine | 0.7592 | -0.39735 | 0.009447 | 0.94151 |
| trans-Aconitate | 0.7872 | -0.34519 | 0.01189 | 0.887388 |
| Guanosine | 1.5342 | 0.617502 | 0.014427 | 0.844777 |
| Choline | 1.5001 | 0.58506 | 0.020593 | 0.728029 |
| Oxalate | 0.7970 | -0.32739 | 0.020917 | 0.75602 |
| Fad | 0.7513 | -0.41253 | 0.022638 | 0.753862 |
| Glutamine | 0.8357 | -0.25901 | 0.026266 | 0.71927 |
| Mannose | 0.8255 | -0.2766 | 0.030791 | 0.678268 |
| Threonine | 0.8386 | -0.25396 | 0.03082 | 0.704187 |
| Aspartate | 0.7507 | -0.4137 | 0.032183 | 0.710225 |
| Adenosine | 1.3494 | 0.432318 | 0.033546 | 0.715683 |
| Adenosine 3',5'-diphosphate | 0.6440 | -0.63497 | 0.033961 | 0.73262 |
| Inosine | 1.6030 | 0.680763 | 0.038461 | 0.699773 |
| Nicotinamide | 1.2965 | 0.374631 | 0.043227 | 0.669243 |
| Glutamate | 0.8461 | -0.2411 | 0.045185 | 0.669304 |
| 3-Hydroxymethylglutarate | 0.8464 | -0.24059 | 0.049068 | 0.651986 |
| Adenine | 0.7791 | -0.36006 | 0.049797 | 0.663311 |

**Supplementary Table 3.**
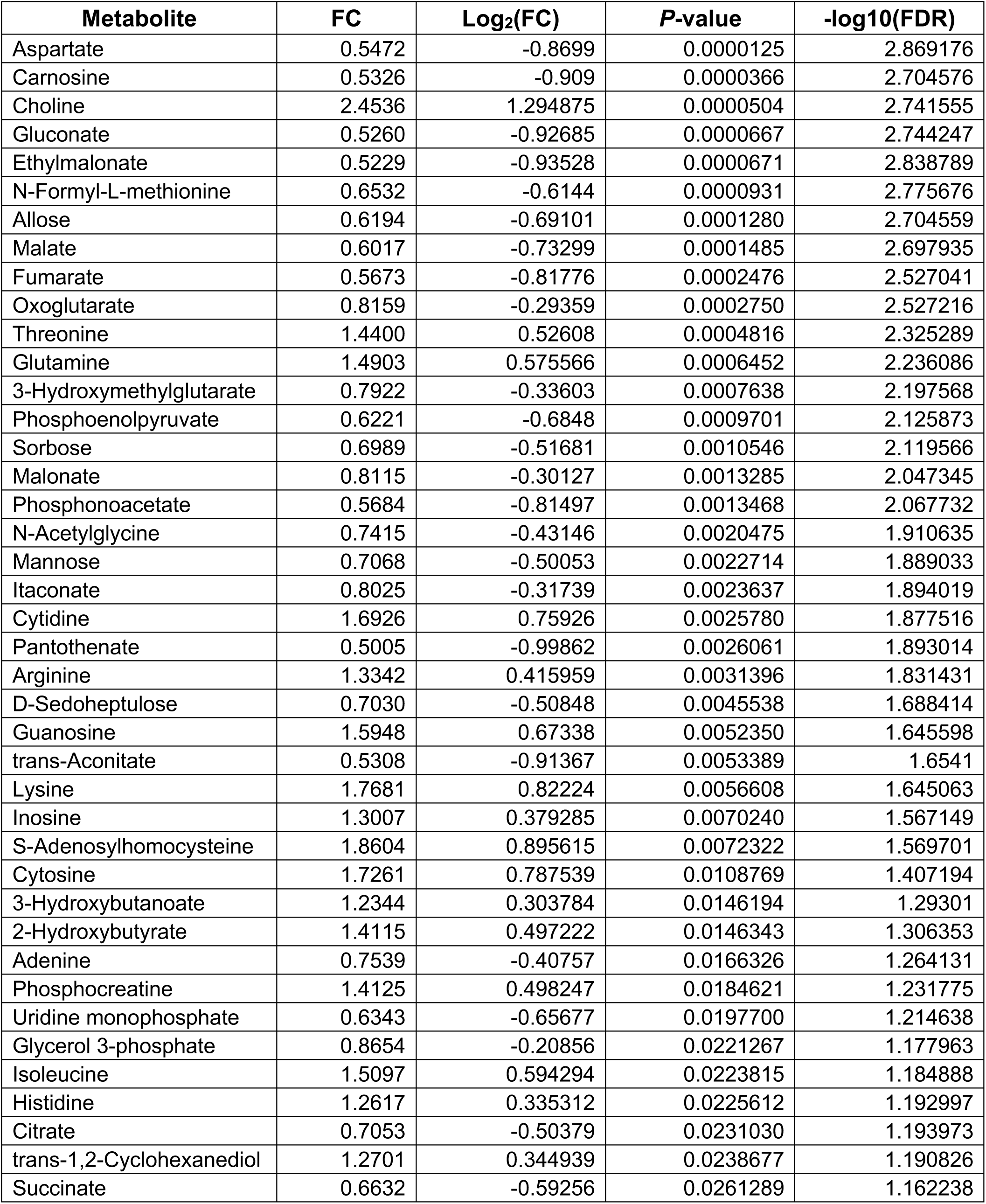
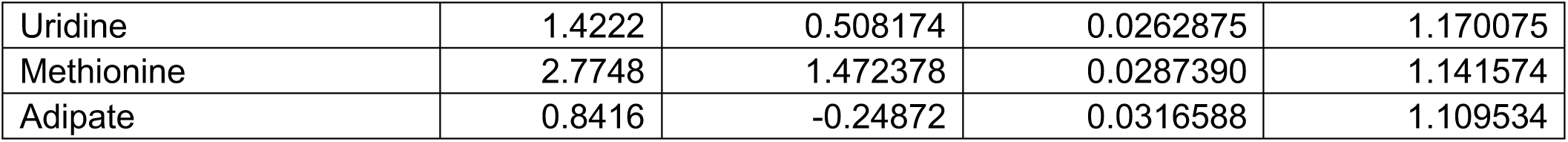
Differential mitochondrial metabolites in control and *Slc25a34* KO primary myotubes under a starved condition (*P-*value < 0.05), related to Figure 3.

**Supplementary Table 4.** Differential whole-cell metabolites in control and *Slc25a34* KO primary myotubes under a starved condition (*P-*value < 0.05), related to Figure S3.

| Metabolite | FC | Log<sub>2</sub>(FC) | P-value | -log<sub>10</sub>(FDR) |
| --- | --- | --- | --- | --- |
| Sorbose | 0.7251 | -0.46377 | 3.19817E-05 | 2.457673 |
| Inosine | 1.4421 | 0.528212 | 3.83719E-05 | 2.67959 |
| gamma-Aminobutyrate | 0.7356 | -0.44298 | 0.000157432 | 2.242603 |
| Methylthioadenosine | 0.7681 | -0.38065 | 0.000172795 | 2.327103 |
| Cytidine monophosphate | 0.8175 | -0.29077 | 0.000244872 | 2.272604 |
| Adenine | 0.7792 | -0.35997 | 0.000344334 | 2.203745 |
| Sorbitol | 0.6689 | -0.58003 | 0.000405724 | 2.199441 |
| Oxalate | 0.8039 | -0.31495 | 0.000411093 | 2.251723 |
| Adipate | 0.4482 | -1.15788 | 0.000417711 | 2.29594 |
| Malonate | 0.8139 | -0.29704 | 0.000568926 | 2.207518 |
| Uridine diphosphate-N-acetylglucosamine | 0.7142 | -0.48565 | 0.000839704 | 2.07984 |
| Histidine | 1.4653 | 0.551209 | 0.001348359 | 1.911949 |
| Betaine | 0.7053 | -0.50369 | 0.001761089 | 1.830736 |
| Sucrose | 0.7713 | -0.3747 | 0.00199262 | 1.809277 |
| beta-Nicotinamide Adenine Dinucleotide | 0.7455 | -0.42374 | 0.002399434 | 1.758556 |
| 4-Guanidinobutanoate | 0.8060 | -0.31119 | 0.002873019 | 1.708355 |
| Phosphocreatine | 0.7618 | -0.39243 | 0.00364187 | 1.631698 |
| N,N,N-Trimethyllysine | 0.8148 | -0.29555 | 0.004890016 | 1.528536 |
| Deoxycarnitine | 0.7641 | -0.38808 | 0.006405057 | 1.434804 |
| Oxoproline | 1.3700 | 0.454197 | 0.006428375 | 1.455502 |
| Pyroglutamate | 1.3700 | 0.454197 | 0.006428375 | 1.476692 |
| CDP-ethanolamine | 0.7998 | -0.32232 | 0.006431727 | 1.496669 |
| Proline | 1.4298 | 0.515806 | 0.007031136 | 1.477276 |
| Taurine | 0.8033 | -0.31606 | 0.011400605 | 1.285857 |
| Choline | 1.5207 | 0.604738 | 0.011741597 | 1.290786 |
| Methionine | 1.1438 | 0.193868 | 0.013801152 | 1.237632 |
| Glutathione oxidized | 1.2156 | 0.28162 | 0.015839088 | 1.194207 |
| Phenylalanine | 1.2943 | 0.372201 | 0.021233736 | 1.082705 |
| L-Carnitine | 0.8329 | -0.26371 | 0.021998666 | 1.082575 |
| Adenosine-monophosphate | 0.7202 | -0.47344 | 0.029065758 | 0.976313 |
| 3-Hydroxymethylglutarate | 0.8475 | -0.23871 | 0.032038752 | 0.94826 |
| Guanosine | 1.1805 | 0.239428 | 0.035450839 | 0.918097 |
| Nicotinamide | 0.8845 | -0.17706 | 0.038795485 | 0.892306 |
| Glutamine | 1.1807 | 0.239645 | 0.049597741 | 0.798591 |

**Supplementary Table 5.**
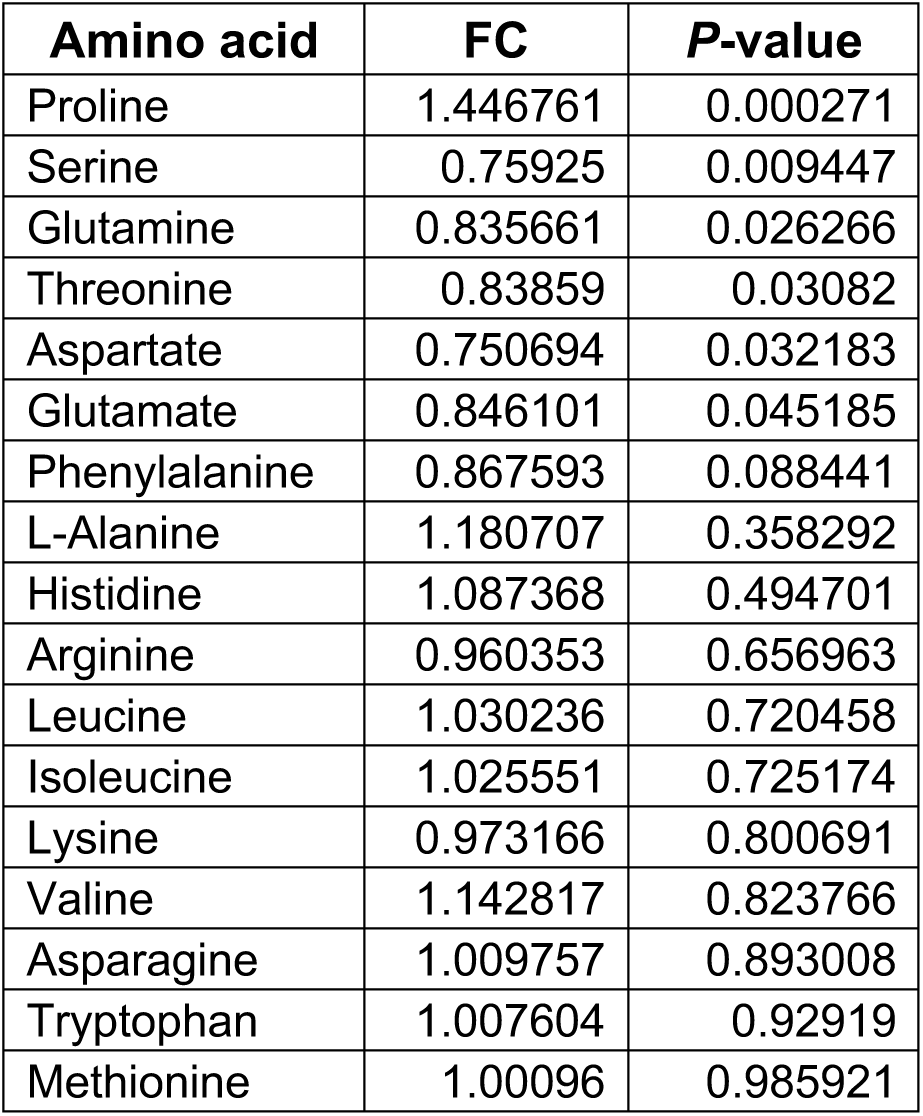
Intracellular amino acid levels in control and *Slc25a34* KO primary myotubes under a fed condition, related to Figure 6.

**Supplementary Table 6.** Intracellular amino acid levels in control and *Slc25a34* KO primary myotubes under a starved condition, related to Figure 6.

| Amino acid | FC | P-value |
| --- | --- | --- |
| Histidine | 1.4653 | 0.001348 |
| Proline | 1.4297 | 0.007031 |
| Methionine | 1.1438 | 0.013801 |
| Phenylalanine | 1.2943 | 0.021234 |
| Glutamine | 1.1807 | 0.049598 |
| Lysine | 1.1699 | 0.081041 |
| Threonine | 1.1914 | 0.083601 |
| Valine | 1.3602 | 0.102727 |
| Serine | 1.1251 | 0.143351 |
| L-Alanine | 0.8758 | 0.310715 |
| Leucine | 1.0543 | 0.337289 |
| Asparagine | 1.0918 | 0.379808 |
| Aspartate | 1.0759 | 0.461471 |
| Isoleucine | 1.0492 | 0.47854 |
| Arginine | 1.0482 | 0.478971 |
| Tryptophan | 1.0321 | 0.646365 |
| Glutamate | 0.9547 | 0.684545 |

**Supplementary Table 7.** Intracellular amino acid levels in control and *Slc25a35* KO primary myotubes under a fed condition, related to Figure 6.

| **Amino acid** | **FC** | ***P*-value** |
| --- | --- | --- |
| Threonine | 1.3529 | 0.01786137 |
| Glutamate | 1.0978 | 0.09411465 |
| Valine | 0.8798 | 0.09809924 |
| Methoinine | 0.8309 | 0.13839674 |
| Aspartate | 1.0577 | 0.18632468 |
| Arginine | 0.8879 | 0.23051777 |
| Serine | 1.0533 | 0.27774215 |
| Glycine | 1.0634 | 0.36862782 |
| Glutamine | 1.0804 | 0.37546956 |
| Asparagine | 1.0489 | 0.55186885 |
| Alanine | 1.0192 | 0.61704747 |
| Leucine | 1.0640 | 0.66619551 |
| Histidine | 0.9805 | 0.67797706 |
| Tryptophan | 1.0451 | 0.7548336 |
| Phenylalanine | 0.9617 | 0.76975901 |
| Lysine | 1.0247 | 0.78947997 |
| Isoleucine | 1.0237 | 0.81831771 |
| Tyrosine | 1.0106 | 0.83731928 |

**Supplementary Table 8.** Intracellular amino acid levels in control and *Slc25a35* KO primary myotubes under a starved condition, related to Figure 6.

| Amino acid | FC | P-value |
| --- | --- | --- |
| Glutamate | 1.2337 | 0.02481925 |
| Phenylalanine | 0.7522 | 0.04059091 |
| Aspartate | 1.1090 | 0.10800255 |
| Arginine | 0.6506 | 0.17986799 |
| Threonine | 0.9215 | 0.33515299 |
| Glutamine | 1.0274 | 0.4784421 |
| Valine | 1.4269 | 0.48237451 |
| Leucine | 1.0665 | 0.54554677 |
| Isoleucine | 1.0586 | 0.64544186 |
| Alanine | 0.9811 | 0.65888572 |
| Serine | 0.9524 | 0.66326534 |
| Methionine | 1.1417 | 0.81411647 |
| Asparagine | 1.0213 | 0.81919316 |
| Histidine | 1.0573 | 0.82182132 |
| Tryptophan | 0.9585 | 0.89250769 |
| Tyrosine | 0.9946 | 0.96015322 |

**Supplementary Table 9.**
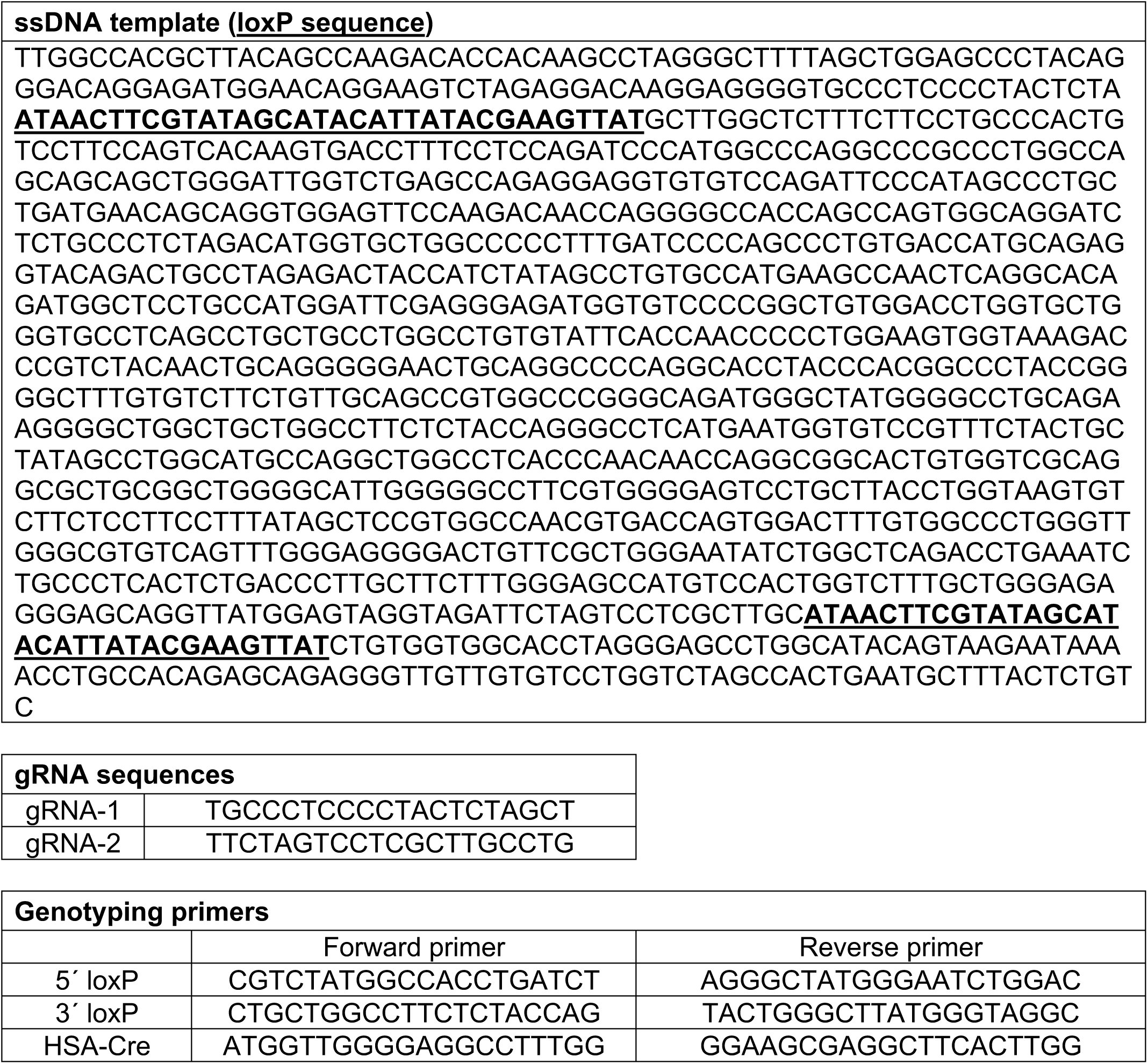
sgRNAs, ssDNA template and genotyping primers for generation of *Slc25a34*^flox/-^ mice.

**Supplementary Table 10.**
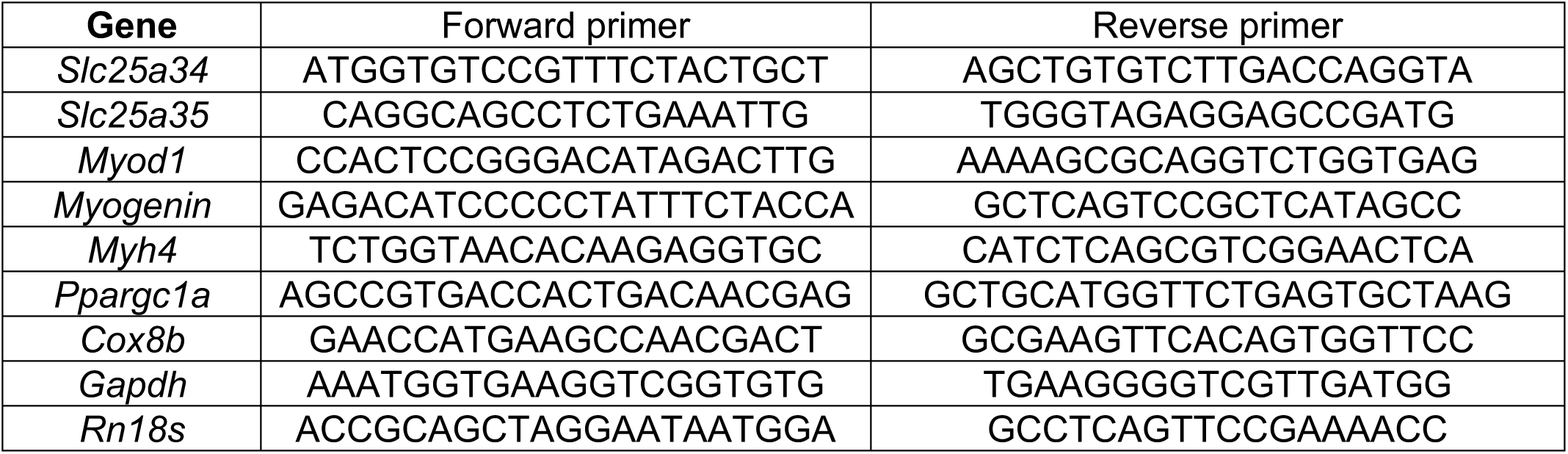
RT-qPCR primer sequences.

## References

1. R. P. Chakrabarty, N. S. Chandel, Beyond ATP, new roles of mitochondria. Biochem (Lond) 44, 2–8 (2022).

2. J. B. Spinelli, M. C. Haigis, The multifaceted contributions of mitochondria to cellular metabolism. Nature cell biology 20, 745–754 (2018).

3. M. Granath-Panelo, S. Kajimura, Mitochondrial heterogeneity and adaptations to cellular needs. Nature cell biology 26, 674–686 (2024).

4. J. Axsom, T. TeSlaa, W. D. Lee, Q. Chu, A. Cowan, M. R. Bornstein, M. D. Neinast, C. R. Bartman, M. C. Blair, K. Li, C. Thorsheim, J. D. Rabinowitz, Z. Arany, Quantification of nutrient fluxes during acute exercise in mice. Cell metabolism 36, 2560–2579 e2565 (2024).

5. G. Park, J. A. Haley, J. Le, S. M. Jung, T. P. Fitzgibbons, E. D. Korobkina, H. Li, S. M. Fluharty, Q. Chen, J. B. Spinelli, C. M. Trivedi, C. Jang, D. A. Guertin, Quantitative analysis of metabolic fluxes in brown fat and skeletal muscle during thermogenesis. Nat Metab 5, 1204–1220 (2023).

6. A. R. P. Verkerke, D. Wang, N. Yoshida, Z. H. Taxin, X. Shi, S. Zheng, Y. Li, C. Auger, S. Oikawa, J. S. Yook, M. Granath-Panelo, W. He, G. F. Zhang, M. Matsushita, M. Saito, R. E. Gerszten, E. L. Mills, A. S. Banks, Y. Ishihama, P. J. White, R. W. McGarrah, T. Yoneshiro, S. Kajimura, BCAA-nitrogen flux in brown fat controls metabolic health independent of thermogenesis. Cell 187, 2359–2374 e2318 (2024).

7. R. R. Wolfe, The underappreciated role of muscle in health and disease. Am J Clin Nutr 84, 475–482 (2006).

8. R. A. Bell, M. Al-Khalaf, L. A. Megeney, The beneficial role of proteolysis in skeletal muscle growth and stress adaptation. Skelet Muscle 6, 16 (2016).

9. J. A. B. Smith, K. A. Murach, K. A. Dyar, J. R. Zierath, Exercise metabolism and adaptation in skeletal muscle. Nature reviews. Molecular cell biology 24, 607–632 (2023).

10. R. L. Smith, M. R. Soeters, R. C. I. Wust, R. H. Houtkooper, Metabolic Flexibility as an Adaptation to Energy Resources and Requirements in Health and Disease. Endocrine reviews 39, 489–517 (2018).

11. G. D. Lopaschuk, Q. G. Karwi, R. Tian, A. R. Wende, E. D. Abel, Cardiac Energy Metabolism in Heart Failure. Circulation research 128, 1487–1513 (2021).

12. H. L. Kornberg, The role and control of the glyoxylate cycle in Escherichia coli. The Biochemical journal 99, 1–11 (1966).

13. O. E. Owen, S. C. Kalhan, R. W. Hanson, The key role of anaplerosis and cataplerosis for citric acid cycle function. The Journal of biological chemistry 277, 30409–30412 (2002).

14. M. Inigo, S. Deja, S. C. Burgess, Ins and Outs of the TCA Cycle: The Central Role of Anaplerosis. Annual review of nutrition 41, 19–47 (2021).

15. P. K. Arnold, L. W. S. Finley, Regulation and function of the mammalian tricarboxylic acid cycle. The Journal of biological chemistry 299, 102838 (2023).

16. K. E. Wellen, G. Hatzivassiliou, U. M. Sachdeva, T. V. Bui, J. R. Cross, C. B. Thompson, ATP-citrate lyase links cellular metabolism to histone acetylation. Science 324, 1076–1080 (2009).

17. G. Hatzivassiliou, F. Zhao, D. E. Bauer, C. Andreadis, A. N. Shaw, D. Dhanak, S. R. Hingorani, D. A. Tuveson, C. B. Thompson, ATP citrate lyase inhibition can suppress tumor cell growth. Cancer Cell 8, 311–321 (2005).

18. C. N. Cunningham, J. Rutter, 20,000 picometers under the OMM: diving into the vastness of mitochondrial metabolite transport. EMBO reports 21, e50071 (2020).

19. D. K. Bricker, E. B. Taylor, J. C. Schell, T. Orsak, A. Boutron, Y. C. Chen, J. E. Cox, C. M. Cardon, J. G. Van Vranken, N. Dephoure, C. Redin, S. Boudina, S. P. Gygi, M. Brivet, C. S. Thummel, J. Rutter, A mitochondrial pyruvate carrier required for pyruvate uptake in yeast, Drosophila, and humans. Science 337, 96–100 (2012).

20. S. Herzig, E. Raemy, S. Montessuit, J. L. Veuthey, N. Zamboni, B. Westermann, E. R. Kunji, J. C. Martinou, Identification and functional expression of the mitochondrial pyruvate carrier. Science 337, 93–96 (2012).

21. G. Fiermonte, L. Palmieri, V. Dolce, F. M. Lasorsa, F. Palmieri, M. J. Runswick, J. E. Walker, The sequence, bacterial expression, and functional reconstitution of the rat mitochondrial dicarboxylate transporter cloned via distant homologs in yeast and Caenorhabditis elegans. The Journal of biological chemistry 273, 24754–24759 (1998).

22. V. Dolce, A. R. Cappello, L. Capobianco, Mitochondrial tricarboxylate and dicarboxylate-tricarboxylate carriers: from animals to plants. IUBMB Life 66, 462–471 (2014).

23. F. Palmieri, M. Monne, Discoveries, metabolic roles and diseases of mitochondrial carriers: A review. Biochimica et biophysica acta 1863, 2362–2378 (2016).

24. J. J. Ruprecht, E. R. S. Kunji, The SLC25 Mitochondrial Carrier Family: Structure and Mechanism. Trends Biochem Sci 45, 244–258 (2020).

25. N. D. Amoedo, G. Punzi, E. Obre, D. Lacombe, A. De Grassi, C. L. Pierri, R. Rossignol, AGC1/2, the mitochondrial aspartate-glutamate carriers. Biochim Biophys Acta 1863, 2394–2412 (2016).

26. M. Monne, D. V. Miniero, V. Iacobazzi, F. Bisaccia, G. Fiermonte, The mitochondrial oxoglutarate carrier: from identification to mechanism. J Bioenerg Biomembr 45, 1–13 (2013).

27. F. Bisaccia, C. Indiveri, F. Palmieri, Purification of reconstitutively active alpha-oxoglutarate carrier from pig heart mitochondria. Biochimica et biophysica acta 810, 362–369 (1985).

28. P. Borst, The malate-aspartate shuttle (Borst cycle): How it started and developed into a major metabolic pathway. IUBMB Life 72, 2241–2259 (2020).

29. C. Wolfrum, Z. Gerhart-Hines, Fueling the fire of adipose thermogenesis. Science 375, 1229–1231 (2022).

30. P. Cohen, S. Kajimura, The cellular and functional complexity of thermogenic fat. Nature reviews. Molecular cell biology 22, 393–409 (2021).

31. T. Yamamuro, D. Katoh, G. M. Silva, H. Nishida, S. Oikawa, Y. Higuchi, D. Wang, M. Fujimoto, N. Yoshida, M. Li, J. Shin, Z. Zhao, J. S. Yook, L. Sun, S. Kajimura, Mitochondrial control of glycerolipid synthesis by a PEP shuttle. Cell 189, 2988–3003 e2923 (2026).

32. S. Yu, S. Meng, M. Xiang, H. Ma, Phosphoenolpyruvate carboxykinase in cell metabolism: Roles and mechanisms beyond gluconeogenesis. Molecular metabolism 53, 101257 (2021).

33. E. Possik, A. Al-Mass, M. L. Peyot, R. Ahmad, F. Al-Mulla, S. R. M. Madiraju, M. Prentki, New Mammalian Glycerol-3-Phosphate Phosphatase: Role in beta-Cell, Liver and Adipocyte Metabolism. Frontiers in endocrinology 12, 706607 (2021).

34. M. J. Merrins, B. E. Corkey, R. G. Kibbey, M. Prentki, Metabolic cycles and signals for insulin secretion. Cell metabolism 34, 947–968 (2022).

35. P. Poursharifi, S. R. M. Madiraju, A. Oppong, S. Kajimura, C. J. Nolan, D. P. Blondin, M. Prentki, Glycerolipid Cycling in Thermogenesis, Energy Homeostasis and Signaling. Physiological reviews 105, 2449–2499 (2025).

36. F. Palmieri, The mitochondrial transporter family SLC25: identification, properties and physiopathology. Mol Aspects Med 34, 465–484 (2013).

37. D. Bertoni, M. Tsenkov, P. Magana, S. Nair, I. Pidruchna, M. Querino Lima Afonso, A. Midlik, U. Paramval, D. Lawal, A. Tanweer, M. Last, R. Patel, A. Laydon, D. Lasecki, N. Dietrich, H. Tomlinson, A. Zidek, T. Green, O. Kovalevskiy, A. Lau, S. Kandathil, N. Bordin, I. Sillitoe, M. Mirdita, D. Jones, C. Orengo, M. Steinegger, J. R. Fleming, S. Velankar, AlphaFold Protein Structure Database 2025: a redesigned interface and updated structural coverage. Nucleic Acids Res 54, D358–D362 (2026).

38. J. Tai, R. M. Guerra, S. W. Rogers, Z. Fang, L. K. Muehlbauer, E. Shishkova, K. A. Overmyer, J. J. Coon, D. J. Pagliarini, Hem25p is required for mitochondrial IPP transport in fungi. Nat Cell Biol 25, 1616–1624 (2023).

39. C. M. Marobbio, G. Giannuzzi, E. Paradies, C. L. Pierri, F. Palmieri, alpha-Isopropylmalate, a leucine biosynthesis intermediate in yeast, is transported by the mitochondrial oxalacetate carrier. J Biol Chem 283, 28445–28453 (2008).

40. C. Auger, H. Nishida, B. Yuan, G. M. Silva, M. Fujimoto, M. Li, D. Katoh, D. Wang, M. Granath-Panelo, J. Shin, R. Witte, J. S. Yook, A. R. P. Verkerke, A. S. Banks, S. Hui, L. Sun, S. Kajimura, Mitochondrial control of fuel switching via carnitine biosynthesis. Science 391, eady5532 (2026).

41. A. Khan, F. S. Yen, G. Unlu, N. L. DelGaudio, R. Erdal, M. Xiao, K. Wangdu, K. Cho, E. R. Gamazon, G. J. Patti, K. Birsoy, Machine-learning-guided discovery of SLC25A45 as a mediator of mitochondrial methylated amino acid import and carnitine synthesis. Cell metabolism 37, 2220–2232 e2228 (2025).

42. M. M. Dias, M. S. King, E. Shokry, S. Lilla, N. Paul, P. Thomason, S. Zanivan, D. Sumpton, E. R. S. Kunji, T. MacVicar, SLC25A45 is required for mitochondrial uptake of methylated amino acids and de novo carnitine biosynthesis. Molecular cell 85, 4093–4104 e4098 (2025).

43. J. Han, S. Gagnon, T. Eckle, C. H. Borchers, Metabolomic analysis of key central carbon metabolism carboxylic acids as their 3-nitrophenylhydrazones by UPLC/ESI-MS. Electrophoresis 34, 2891–2900 (2013).

44. M. S. King, M. Kerr, P. G. Crichton, R. Springett, E. R. S. Kunji, Formation of a cytoplasmic salt bridge network in the matrix state is a fundamental step in the transport mechanism of the mitochondrial ADP/ATP carrier. Biochim Biophys Acta 1857, 14–22 (2016).

45. S. Schmid, B. Heim-Kupr, J. Perez-Schindler, S. Mansingh, M. Beer, N. Mittal, N. Ehrenfeuchter, C. Handschin, PGC-1beta modulates catabolism and fiber atrophy in the fasting-response of specific skeletal muscle beds. Mol Metab 66, 101643 (2022).

46. K. Ramachandran, M. D. Senagolage, M. A. Sommars, C. R. Futtner, Y. Omura, A. L. Allred, G. D. Barish, Dynamic enhancers control skeletal muscle identity and reprogramming. PloS Biol 17, e3000467 (2019).

47. J. Lin, C. Handschin, B. M. Spiegelman, Metabolic control through the PGC-1 family of transcription coactivators. Cell metabolism 1, 361–370 (2005).

48. G. D. Barish, V. A. Narkar, R. M. Evans, PPAR delta: a dagger in the heart of the metabolic syndrome. The Journal of clinical investigation 116, 590–597 (2006).

49. G. R. Steinberg, D. G. Hardie, New insights into activation and function of the AMPK. Nat Rev Mol Cell Biol 24, 255–272 (2023).

50. M. Baresic, S. Salatino, B. Kupr, E. van Nimwegen, C. Handschin, Transcriptional network analysis in muscle reveals AP-1 as a partner of PGC-1alpha in the regulation of the hypoxic gene program. Mol Cell Biol 34, 2996–3012 (2014).

51. R. Furrer, B. Heim, S. Schmid, S. Dilbaz, V. Adak, K. J. V. Nordstrom, D. Ritz, S. A. Steurer, J. Walter, C. Handschin, Molecular control of endurance training adaptation in male mouse skeletal muscle. Nat Metab 5, 2020–2035 (2023).

52. H. Miura, R. M. Quadros, C. B. Gurumurthy, M. Ohtsuka, Easi-CRISPR for creating knock-in and conditional knockout mouse models using long ssDNA donors. Nat Protoc 13, 195–215 (2018).

53. W. W. Chen, E. Freinkman, D. M. Sabatini, Rapid immunopurification of mitochondria for metabolite profiling and absolute quantification of matrix metabolites. Nature protocols 12, 2215–2231 (2017).

54. C. A. Bellissimo, M. C. Garibotti, C. G. R. Perry, Mitochondrial stress responses in Duchenne muscular dystrophy: metabolic dysfunction or adaptive reprogramming? American journal of physiology. Cell physiology 323, C718–C730 (2022).

55. U. Sharma, S. Atri, M. C. Sharma, C. Sarkar, N. R. Jagannathan, Skeletal muscle metabolism in Duchenne muscular dystrophy (DMD): an in-vitro proton NMR spectroscopy study. Magn Reson Imaging 21, 145–153 (2003).

56. S. Dadgar, Z. Wang, H. Johnston, A. Kesari, K. Nagaraju, Y. W. Chen, D. A. Hill, T. A. Partridge, M. Giri, R. J. Freishtat, J. Nazarian, J. Xuan, Y. Wang, E. P. Hoffman, Asynchronous remodeling is a driver of failed regeneration in Duchenne muscular dystrophy. J Cell Biol 207, 139–158 (2014).

57. D. R. Lynch, G. Farmer, Mitochondrial and metabolic dysfunction in Friedreich ataxia: update on pathophysiological relevance and clinical interventions. Neuronal Signal 5, NS20200093 (2021).

58. E. Indelicato, K. Faserl, M. Amprosi, W. Nachbauer, R. Schneider, J. Wanschitz, B. Sarg, S. Boesch, Skeletal muscle proteome analysis underpins multifaceted mitochondrial dysfunction in Friedreich’s ataxia. Front Neurosci 17, 1289027 (2023).

59. E. Indelicato, A. Kirchmair, M. Amprosi, S. Steixner, W. Nachbauer, A. Eigentler, N. Wahl, G. Apostolova, A. Krogsdam, R. Schneider, J. Wanschitz, Z. Trajanoski, S. Boesch, Skeletal muscle transcriptomics dissects the pathogenesis of Friedreich’s ataxia. Hum Mol Genet 32, 2241–2250 (2023).

60. K. A. Kras, N. Hoffman, L. R. Roust, S. H. Patel, C. C. Carroll, C. S. Katsanos, Plasma Amino Acids Stimulate Uncoupled Respiration of Muscle Subsarcolemmal Mitochondria in Lean but Not Obese Humans. J Clin Endocrinol Metab 102, 4515–4525 (2017).

61. M. E. Bizeau, W. T. Willis, J. R. Hazel, Differential responses to endurance training in subsarcolemmal and intermyofibrillar mitochondria. J Appl Physiol (1985) 85, 1279–1284 (1998).

62. G. Y. Liu, D. M. Sabatini, mTOR at the nexus of nutrition, growth, ageing and disease. Nat Rev Mol Cell Biol 21, 183–203 (2020).

63. C. A. Goodman, D. M. Mabrey, J. W. Frey, M. H. Miu, E. K. Schmidt, P. Pierre, T. A. Hornberger, Novel insights into the regulation of skeletal muscle protein synthesis as revealed by a new nonradioactive in vivo technique. FASEB J 25, 1028–1039 (2011).

64. R. A. Saxton, D. M. Sabatini, mTOR Signaling in Growth, Metabolism, and Disease. Cell 168, 960–976 (2017).

65. N. S. Chandel, Signaling and Metabolism. Cold Spring Harbor perspectives in biology 13, a040600 (2021).

66. E. Ostergaard, F. J. Hansen, N. Sorensen, M. Duno, J. Vissing, P. L. Larsen, O. Faeroe, S. Thorgrimsson, F. Wibrand, E. Christensen, M. Schwartz, Mitochondrial encephalomyopathy with elevated methylmalonic acid is caused by SUCLA2 mutations. Brain 130, 853–861 (2007).

67. R. Carrozzo, C. Dionisi-Vici, U. Steuerwald, S. Lucioli, F. Deodato, S. Di Giandomenico, E. Bertini, B. Franke, L. A. Kluijtmans, M. C. Meschini, C. Rizzo, F. Piemonte, R. Rodenburg, R. Santer, F. M. Santorelli, A. van Rooij, D. Vermunt-de Koning, E. Morava, R. A. Wevers, SUCLA2 mutations are associated with mild methylmalonic aciduria, Leigh-like encephalomyopathy, dystonia and deafness. Brain 130, 862–874 (2007).

68. R. S. Kaplan, J. A. Mayor, N. Johnston, D. L. Oliveira, Purification and characterization of the reconstitutively active tricarboxylate transporter from rat liver mitochondria. J Biol Chem 265, 13379–13385 (1990).

69. S. Balaraju, A. Topf, G. McMacken, V. P. Kumar, A. Pechmann, H. Roper, S. Vengalil, K. Polavarapu, S. Nashi, N. P. Mahajan, I. A. Barbosa, C. Deshpande, R. W. Taylor, J. Cossins, D. Beeson, S. Laurie, J. Kirschner, R. Horvath, R. McFarland, A. Nalini, H. Lochmuller, Congenital myasthenic syndrome with mild intellectual disability caused by a recurrent SLC25A1 variant. Eur J Hum Genet 28, 373–377 (2020).

70. A. Chaouch, V. Porcelli, D. Cox, S. Edvardson, P. Scarcia, A. De Grassi, C. L. Pierri, J. Cossins, S. H. Laval, H. Griffin, J. S. Muller, T. Evangelista, A. Topf, A. Abicht, A. Huebner, M. von der Hagen, K. Bushby, V. Straub, R. Horvath, O. Elpeleg, J. Palace, J. Senderek, D. Beeson, L. Palmieri, H. Lochmuller, Mutations in the Mitochondrial Citrate Carrier SLC25A1 are Associated with Impaired Neuromuscular Transmission. J Neuromuscul Dis 1, 75–90 (2014).

71. G. Punzi, V. Porcelli, M. Ruggiu, M. F. Hossain, A. Menga, P. Scarcia, A. Castegna, R. Gorgoglione, C. L. Pierri, L. Laera, F. M. Lasorsa, E. Paradies, I. Pisano, C. M. T. Marobbio, E. Lamantea, D. Ghezzi, V. Tiranti, S. Giannattasio, M. A. Donati, R. Guerrini, L. Palmieri, F. Palmieri, A. De Grassi, SLC25A10 biallelic mutations in intractable epileptic encephalopathy with complex I deficiency. Hum Mol Genet 27, 499–504 (2018).

72. X. Gu, J. M. Orozco, R. A. Saxton, K. J. Condon, G. Y. Liu, P. A. Krawczyk, S. M. Scaria, J. W. Harper, S. P. Gygi, D. M. Sabatini, SAMTOR is an S-adenosylmethionine sensor for the mTORC1 pathway. Science 358, 813–818 (2017).

73. J. L. Jewell, Y. C. Kim, R. C. Russell, F. X. Yu, H. W. Park, S. W. Plouffe, V. S. Tagliabracci, K. L. Guan, Metabolism. Differential regulation of mTORC1 by leucine and glutamine. Science 347, 194–198 (2015).

74. L. Begum, M. A. Jalil, K. Kobayashi, M. Iijima, M. X. Li, T. Yasuda, M. Horiuchi, A. del Arco, J. Satrustegui, T. Saheki, Expression of three mitochondrial solute carriers, citrin, aralar1 and ornithine transporter, in relation to urea cycle in mice. Biochim Biophys Acta 1574, 283–292 (2002).

75. K. Kobayashi, D. S. Sinasac, M. Iijima, A. P. Boright, L. Begum, J. R. Lee, T. Yasuda, S. Ikeda, R. Hirano, H. Terazono, M. A. Crackower, I. Kondo, L. C. Tsui, S. W. Scherer, T. Saheki, The gene mutated in adult-onset type II citrullinaemia encodes a putative mitochondrial carrier protein. Nat Genet 22, 159–163 (1999).

76. R. Wibom, F. M. Lasorsa, V. Tohonen, M. Barbaro, F. H. Sterky, T. Kucinski, K. Naess, M. Jonsson, C. L. Pierri, F. Palmieri, A. Wedell, AGC1 deficiency associated with global cerebral hypomyelination. N Engl J Med 361, 489–495 (2009).

77. J. E. Kokoszka, K. G. Waymire, A. Flierl, K. M. Sweeney, A. Angelin, G. R. MacGregor, D. C. Wallace, Deficiency in the mouse mitochondrial adenine nucleotide translocator isoform 2 gene is associated with cardiac noncompaction. Biochim Biophys Acta 1857, 1203–1212 (2016).

78. B. H. Graham, K. G. Waymire, B. Cottrell, I. A. Trounce, G. R. MacGregor, D. C. Wallace, A mouse model for mitochondrial myopathy and cardiomyopathy resulting from a deficiency in the heart/muscle isoform of the adenine nucleotide translocator. Nat Genet 16, 226–234 (1997).

79. L. Palmieri, S. Alberio, I. Pisano, T. Lodi, M. Meznaric-Petrusa, J. Zidar, A. Santoro, P. Scarcia, F. Fontanesi, E. Lamantea, I. Ferrero, M. Zeviani, Complete loss-of-function of the heart/muscle-specific adenine nucleotide translocator is associated with mitochondrial myopathy and cardiomyopathy. Hum Mol Genet 14, 3079–3088 (2005).

80. T. D. Goddard, C. C. Huang, E. C. Meng, E. F. Pettersen, G. S. Couch, J. H. Morris, T. E. Ferrin, UCSF ChimeraX: Meeting modern challenges in visualization and analysis. Protein Sci 27, 14–25 (2018).

81. P. Rice, I. Longden, A. Bleasby, EMBOSS: the European Molecular Biology Open Software Suite. Trends Genet 16, 276–277 (2000).

82. J. Mifsud, S. Ravaud, E. M. Krammer, C. Chipot, E. R. Kunji, E. Pebay-Peyroula, F. Dehez, The substrate specificity of the human ADP/ATP carrier AAC1. Mol Membr Biol 30, 160–168 (2013).

83. S. Ravaud, A. Bidon-Chanal, I. Blesneac, P. Machillot, C. Juillan-Binard, F. Dehez, C. Chipot, E. Pebay-Peyroula, Impaired transport of nucleotides in a mitochondrial carrier explains severe human genetic diseases. ACS Chem Biol 7, 1164–1169 (2012).

84. B. Miroux, J. E. Walker, Over-production of proteins in Escherichia coli: mutant hosts that allow synthesis of some membrane proteins and globular proteins at high levels. J Mol Biol 260, 289–298 (1996).

85. L. Dumon-Seignovert, G. Cariot, L. Vuillard, The toxicity of recombinant proteins in Escherichia coli: a comparison of overexpression in BL21(DE3), C41(DE3), and C43(DE3). Protein Expr Purif **37**, 203–206 (2004).

86. E. van Rooij, D. Quiat, B. A. Johnson, L. B. Sutherland, X. Qi, J. A. Richardson, R. J. Kelm, Jr., E. N. Olson, A family of microRNAs encoded by myosin genes governs myosin expression and muscle performance. Dev Cell 17, 662–673 (2009).

87. M. V. Kuleshov, M. R. Jones, A. D. Rouillard, N. F. Fernandez, Q. Duan, Z. Wang, S. Koplev, S. L. Jenkins, K. M. Jagodnik, A. Lachmann, M. G. McDermott, C. D. Monteiro, G. W. Gundersen, A. Ma’ayan, Enrichr: a comprehensive gene set enrichment analysis web server 2016 update. Nucleic Acids Res 44, W90–97 (2016).

